# Cardiac Molecular Analysis Reveals Aging-Associated Metabolic Alterations Promoting Glycosaminoglycans Accumulation Via Hexosamine Biosynthetic Pathway

**DOI:** 10.1101/2023.11.17.567640

**Authors:** Luís F. Grilo, Kip D. Zimmerman, Sobha Puppala, Jeannie Chan, Hillary F. Huber, Ge Li, Avinash Y. L. Jadhav, Benlian Wang, Cun Li, Geoffrey D. Clarke, Thomas C. Register, Paulo J. Oliveira, Peter W. Nathanielsz, Michael Olivier, Susana P. Pereira, Laura A. Cox

## Abstract

Age is a prominent risk factor for cardiometabolic disease, and often leads to heart structural and functional changes. However, precise molecular mechanisms underlying cardiac remodeling and dysfunction resulting from physiological aging per se remain elusive. Understanding these mechanisms requires biological models with optimal translation to humans. Previous research demonstrated that baboons undergo age-related reduction in ejection fraction and increased heart sphericity, mirroring changes observed in humans.

The goal of this study was to identify early cardiac molecular alterations that precede functional adaptations, shedding light on the regulation of age-associated changes.

We performed unbiased transcriptomics of left ventricle (LV) samples from female baboons aged 7.5-22.1 years (human equivalent ∼30-88 years). Weighted-gene correlation network and pathway enrichment analyses were performed to identify potential age-associated mechanisms in LV, with histological validation. Myocardial modules of transcripts negatively associated with age were primarily enriched for cardiac metabolism, including oxidative phosphorylation, tricarboxylic acid cycle, glycolysis, and fatty-acid β-oxidation. Transcripts positively correlated with age suggest upregulation of glucose uptake, pentose phosphate pathway, and hexosamine biosynthetic pathway (HBP), indicating a metabolic shift towards glucose-dependent anabolic pathways. Upregulation of HBP commonly results in increased glycosaminoglycan precursor synthesis. Transcripts involved in glycosaminoglycan synthesis, modification, and intermediate metabolism were also upregulated in older animals, while glycosaminoglycan degradation transcripts were downregulated with age. These alterations would promote glycosaminoglycan accumulation, which was verified histologically. Upregulation of extracellular matrix (ECM)-induced signaling pathways temporally coincided with glycosaminoglycan accumulation. We found a subsequent upregulation of cardiac hypertrophy-related pathways and an increase in cardiomyocyte width.

Overall, our findings revealed a transcriptional shift in metabolism from catabolic to anabolic pathways that leads to ECM glycosaminoglycan accumulation through HBP prior to upregulation of transcripts of cardiac hypertrophy-related pathways. This study illuminates cellular mechanisms that precede development of cardiac hypertrophy, providing novel potential targets to remediate age-related cardiac diseases.

## Introduction

Populations of older individuals have been increasing worldwide, and by 2050, up to one-third of the population in developed countries will be 60 years or older [1]. Age is recognized as a major risk factor for cardiovascular disease (CVD) including coronary heart disease, stroke, and heart failure [2,3]. Aging leads to CVD by promoting heart structural and functional alterations, contributing to cardiac dysfunction [2]. In the European Union, 49 million people were living with CVD in 2019, and it is estimated that by 2030, 130 million adults in the United States will be affected by CVD [4,5].

The heart demands substantial chemical energy, primarily adenosine triphosphate (ATP) production, to drive contraction. Cardiac metabolic flexibility stems from the tight regulation of enzymes, ion channels, and contractile, structural, and membrane proteins, enabling efficient energy conversion in response to stress [6,7]. ATP production in healthy adult myocytes is predominantly derived from fatty acid (FA) oxidation (70-90%) and the remaining from glucose, ketone bodies, and amino acid oxidation [8]. Aging significantly impacts cardiac metabolism, partially through the loss of flexibility on substrate usage [9]. Cardiac mitochondrial electron transport chain gene transcription and efficiency decrease with aging, detrimentally impacting ATP production [6,10]. As age increases, the cardiac use of FA oxidation decreases, and the dependence on and relative rate of glucose metabolism increases [11,12]. Glucose can be channeled to multiple metabolic pathways after uptake in the heart, including glycolysis, glycogen synthesis, pentose phosphate pathway, and hexosamine biosynthetic pathway (HBP) [13].

The decrease in FA import through CD36, leading to a metabolic shift towards increased glucose utilization, hastens the transition from compensated hypertrophy (characterized by enhanced cardiac hypertrophy and interstitial fibrosis) to heart failure [8], while the preservation of FA oxidation prevents the metabolic shift and cardiac hypertrophy development [14]. The metabolic shift and the associated alterations in gene expression, metabolite signaling, and the utilization of glucose-derived carbon for anabolic pathways play a crucial role in the physiological growth of the heart. Conversely, cardiac metabolic inefficiency and disrupted coordination of anabolic activity are now recognized as proximal factors contributing to pathological remodeling [15].

Age-related alterations in cardiac metabolism may underlie the increased susceptibility to CVD in older adults. Despite their potential significance, the implications of this natural age-related metabolic shift in cardiac physiology and its potential contribution to CVD remain largely unexplored. Decreased myocardial reserve capacity, cardiac hypertrophy, and fibrosis are hallmarks of cardiac aging [16]. Nevertheless, the mechanisms that relate the metabolic shift to the emergence of cardiac hypertrophy are not clear. In a previous study, we observed a longitudinal increase in apical pericardial fat, indicating potential metabolic changes, in aging female baboons with no pre-existing health conditions [17]. We also observed compromised left and right ventricle function with aging, suggesting that even healthy physiological aging leads to cardiac remodeling by itself [18]. These biventricular adaptations include a diminished normalized average filling rate, cardiac index, and ejection fraction [18,19]. In addition, LV end-diastolic sphericity index increased while right ventricle sphericity indices dropped with age, suggesting ventricle-specific remodeling [19,20]. LV mass increased with age in males but not females, after correction for body weight-based body surface area estimates, females but not males showed an age-related loss in normalized LV mass [18], highlighting sexual dimorphism and a notable gap in our understanding of the intricate cardiac molecular mechanisms driven by physiological aging in a sex-specific manner.

During cardiac remodeling, the heart undergoes extensive extracellular matrix (ECM) adaptation. Accumulation of ECM is regulated by the balance of degradation and synthesis of its constituents [16,21]. The ECM includes fibrous and non-fibrous components (*e.g.*, proteoglycan and glycosaminoglycan (GAG)) [22]. GAGs are polysaccharide chains that may be secreted upon synthesis (Hyaluronic Acid (HA)) or covalently synthesized on core proteins to form proteoglycans (PG). GAGs can differ in carbohydrate composition, length, and/or sulfation patterns (Chondroitin/Dermatan Sulfate (CS/DS), Heparin/Heparan Sulfate (HS), and Keratan Sulfate (KS)) [23]. PG/GAGs play roles in multiple physiological functions, extending beyond ECM formation. PG/GAGs participate in tissue development and cell signaling through interactions with ECM components, morphogens, cytokines, and growth factors and influence the physical characteristics of tissues [24,25]. Altered PG/GAG structure and function have been linked to heart disease because of their role in cardiac remodeling, hypertrophy, and age-related degeneration of the myocardium [23]. Accumulation of GAGs was previously reported in failing pediatric and adult hearts and in chronic stress [22,26]. However, the aging-related cellular mechanisms that lead to ECM remodeling and to cardiac hypertrophy, and CVD are not well understood.

Initial studies of aging-related cardiac transcriptomics use a traditional comparison of two time points (young vs. old animals) [10,12,27] or more than two specific time points in rodents [28]. The translatability of murine models to understand humans cardiac aging transcriptomics is very limited due to significant developmental, structural and functional cardiovascular differences, including oxygen consumption, adrenergic receptor ratios, and heart rate [29]. Furthermore, the use of senescence-induced animal models lacks reliability due to the non-physiological and uncoordinated promotion of aging mechanisms [28]. While the use of rodent models represents a significant limitation to human translation, the temporal analysis based on classes also compromises the analytical power, causality testing, and effect evaluation. Employing a temporal linear approach increases our understanding of cardiac aging by revealing that not all gene expression consistently follow a linear profile throughout the lifespan [30]. Therefore, our study used a highly translational model, the baboon, a nonhuman primate (NHP) with highly similar genetics that closely mimics human physiology. Cardiac morphology and function have been well characterized in this NHP. The study of healthy cardiac tissue with a short post-mortem interval cannot be readily conducted in humans, highlighting the relevance of using NHP to unravel cardiac aging-specific changes independent from CVD, which is highly prevalent in older individuals [31]. We used age as a continuous variable for a detailed analysis of transcriptional changes during lifespan. Furthermore, it is well-established that sex greatly impacts heart metabolism and response to metabolic diseases. For instance, female hearts use more FAs while male hearts metabolize more glucose [32,33]. Nevertheless, it is worth noting that previous cardiac metabolism studies, both in humans and animals, were predominantly conducted in males [6] so there is a scarcity of research and information specifically focusing on female hearts in this context.

In the present work, we studied 35 healthy female baboons 7.5-22.1 years old (human equivalent ∼30-88 years) [34], to investigate how age-related functional modulation is related to altered cardiac gene expression and consequent metabolic and ECM changes.

For the first time, we identified transcriptional signatures consistent with early metabolic adaptations that lead to an increased flux of the HBP, providing the precursors for GAG synthesis. We found age-related GAG accumulation in the ECM that is temporally coincident with increased expression of genes in ECM-associated pathways. Importantly, these events precede the dysregulation of genes in cardiac hypertrophy-related pathways, suggesting that the cardiac metabolic shift and ECM-GAG accumulation are early mechanisms that prime the heart to later cardiac dysfunction. Therefore, they represent potential diagnostic and therapeutic targets for preventing age-related cardiovascular abnormalities and disorders, including cardiac hypertrophy and subsequent CVD.

## Material and Methods

### Animal model and tissue collection

Thirty-five female baboons (*Papio hamadryas* spp., crosses of olive, hamadryas, and yellow baboons) between 7.5 and 22.1 years old (Figure 1A), with a median age of 13.8, were housed in outdoor social groups composed of 3 to 19 animals at the Southwest National Primate Research Center (SNPRC) at Texas Biomedical Research Institute (TX Biomed), in San Antonio, Texas. The animals were fed normal monkey chow diet low in fat and cholesterol with protein primarily from plant-based sources (Monkey Diet 5LEO, Purina, St Louis, MO, USA) with *ad libitum* availability throughout their lifespan. Multiple lixits provided continuous water availability. Complete veterinary care was provided by SNPRC veterinary staff to all animals throughout their lives.

**Figure 1.**
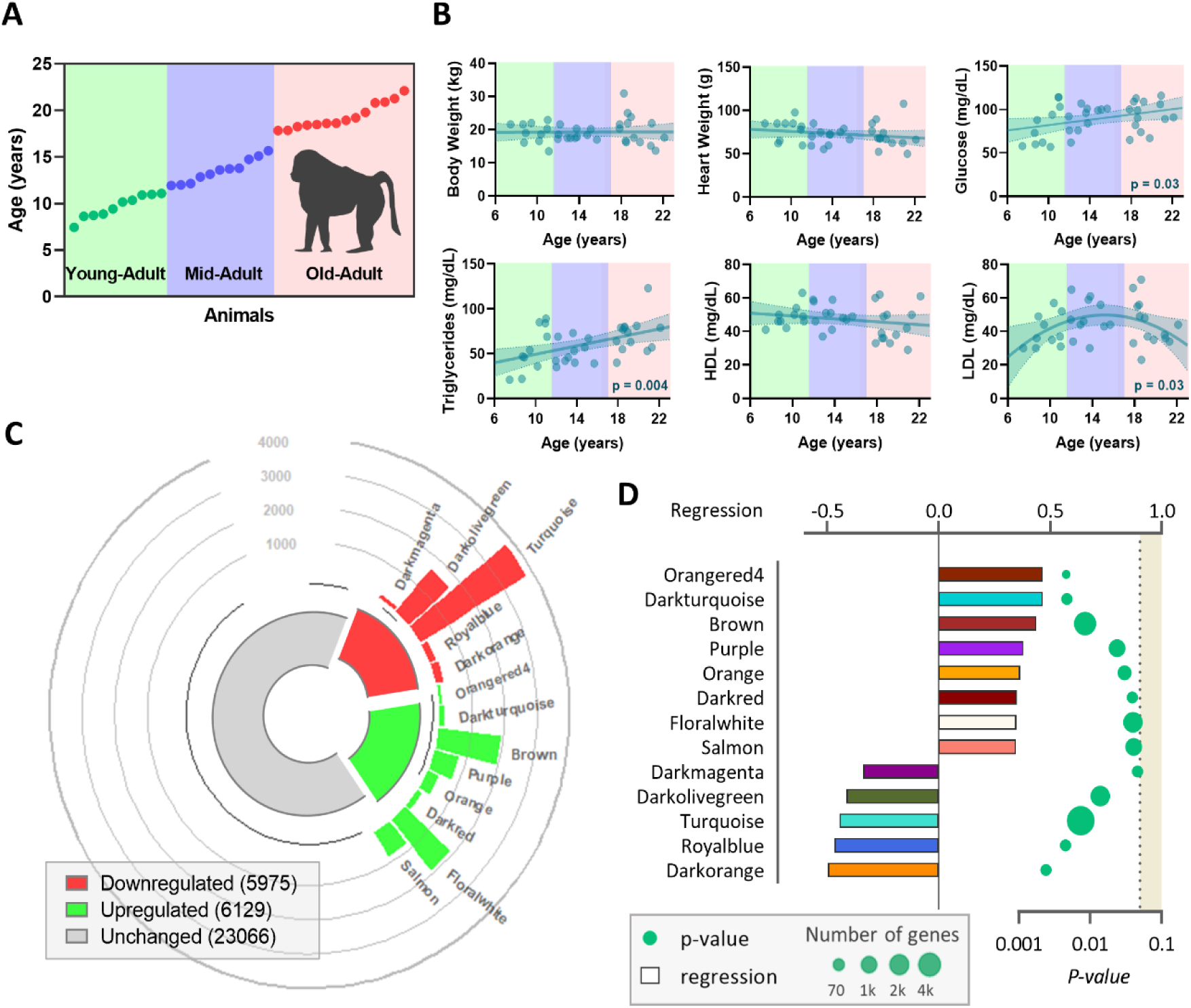
Aging effects on baboon morphometrics, blood clinical data, and cardiac transcriptomics. (A) Schematic representation of the experimental design. (B) Variation of morphometric (body and heart weight) and blood clinical (glucose, triglycerides, HDL, and LDL) measurements with age. (C) Modules positively, in red, and negatively, in green, correlated with age and number of transcripts in each module. The size of the bar is proportional to the number of transcripts from each module. (D) Modules that significantly correlate with age. Each bar is the color of the module and the size represents the regression (top axis), and the dots indicate the module’s p-value (bottom axis) and their size represents the number of transcripts from the respective module.

Before tissue collection, baboons were medicated with ketamine hydrochloride (10 mg/kg intramuscular) and anesthetized using isoflurane (2%) as previously reported [35]. Variations from circadian rhythms were minimized by conducting all collections between 8:00 AM – 10:00 AM. Under general anesthesia, baboons were exsanguinated as approved by the American Veterinary Medical Association. Following cardiac asystole, heart LV tissue was collected, part was snap-frozen in liquid nitrogen, and stored at −80°C while another part was fixed for 48 hours with 4% paraformaldehyde solution, dehydrated, and blocked in paraffin [36].

All procedures were performed in facilities approved by the Association for Assessment and Accreditation of Laboratory Animal Care and previously approved by the TX Biomed Animal Care and Use Committee. TX Biomed animal programs operate according to all U.S. Department of Agriculture and National Institutes of Health (NIH) guidelines and are directed by board-certified veterinarians (DVM). SNPRC veterinarians made all decisions related to animal care.

### Morphometric and blood measurements

Morphometric measures were obtained before animal necropsy and under sedation using anatomical landmarks [37,38]. Blood samples were collected from the femoral vein between 8:00 AM and 10:00 AM and within 5 minutes of intramuscular administration of ketamine in overnight fasted animals.

Plasma lipids (total cholesterol, low-density lipoprotein (LDL) cholesterol, high-density lipoprotein (HDL) cholesterol, and triglyceride) and glucose concentrations were measured on the ACE AXCEL autoanalyzer (Alfa Wasserman Diagnostic Technologies, West Caldwell, NJ, USA) by the Wake Forest Comparative Medicine Clinical Chemistry and Endocrinology Laboratory. Controls from the Centers for Disease Control and Prevention/National Institutes of Health Lipid Standardization Program (Solomon Park, Burien, WA, USA) were used to calibrate and standardize plasma lipids. The assay precision was determined using 5 replicates of pooled samples in each assay.

### RNA isolation from heart samples

Approximately 5 mg of frozen heart LV tissue was homogenized using a BeadBeater (BioSpec, Bartlesville, OK, USA) with zirconia/silica beads in 1 mL RLT buffer (Qiagen, Germantown, MD, USA). RNA was extracted using the Zymo Direct-zol RNA Miniprep Plus kit as recommended by the manufacturer. The quality of the RNA was measured using the TapeStation high-sensitivity RNA ScreenTape (Agilent) and the RNA was stored at −80 °C until further use.

### Gene expression profiling (RNA sequencing)

cDNA libraries were generated from the RNA samples using the Kapa RNA HyperPrep Kit with RiboErase (Roche, Indianapolis, IN, USA). Quality was measured by Agilent D1000 Screen Tape (Santa Clara, CA, USA) according to the manufacturer’s instructions. cDNA libraries were pooled and sequenced using the Illumina Flow cell v 1.5 reagent kit (San Diego, CA, USA) for 2×150 paired-end reads on a NovaSeq 6000 Sequencer.

Before performing alignment, low-quality bases with Phred scores less than 30 were removed. The trimmed reads were aligned using HISAT2 [39] and the olive baboon genome as a reference and then quantified with an expectation-maximization algorithm [40] with Panu_3.0 annotation (NCBI release 103). The criteria to quantify paired-end reads were 100% overlap between reads and the annotated transcripts, and the skipped regions of junction reads must match the transcripts’ introns. Transcripts were normalized using the trimmed mean of the M values method [41]. Normalized read counts in which the sum of all samples was less than 30 counts were filtered resulting in 35,170 transcripts that passed all quality filters.

### Weighted gene co-expression network analysis

Weighted gene co-expression network analysis (WGCNA) was used to identify co-correlated modules of genes. WGCNA (1.70-3) was implemented in R (RStudio version 4.0.5) using the recommended default parameters [42]. Briefly, a correlation matrix with all pair-wise genes was generated and filtered using a soft threshold, power=12, to construct the adjacent matrix. To assess relative gene interconnectedness and proximity, the adjacent matrix was transformed into a topological overlap matrix which allows the identification of gene co-expression modules based on the hierarchical clustering. The parameters used were deep-split value=2; merging threshold=0.25; minimum module size=30. Using age as a continuous correlation trait, module eigengenes were considered significant when correlation≥0.30 and p-value≤0.05 using Pearson correlation.

### Pathway and network enrichment analysis

For annotation of genes by pathway and network enrichment analyses, the directionality of each gene was determined by the module correlation and denoted by a fold-change of −2 for negative correlations and a fold-change of 2 for positive correlations (for quadratic fits, the directionally after the vertex was considered), and the p-value of each gene was based on the p-value of the respective module. Transcript IDs were converted to Gene IDs. For the core analysis, Ingenuity Pathway Analysis (IPA) software (Qiagen) was used. Each canonical pathway p-value was calculated using the two-tailed Fisher’s exact test with p-value ≤ 0.05 considered significant. The activation z-score was used to predict pathway directionality and calculated taking into consideration the impact of the transcript’s directionality on the pathway (*x*_*i*_), the number of transcripts identified in the pathway (*N*), and using the equation:

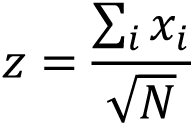

For pathways relevant to cardiac aging that were not included in IPA, manually curated canonical pathways were designed according to the literature reports [24,25,43] and a similar statistical analysis was performed.

### Histology

Three 5 µm sections taken at 150 µm intervals were used for analysis from each animal and were stained with hematoxylin and eosin using the reagents provided in Oil Red O Stain Kit (Abcam, Waltham, MA, USA) for hematoxylin and eosin purchased from StatLab (StatLab, McKinney, TX, USA), and Alcian Blue Stain using Alcian Blue Stain Kit (Abcam, Waltham, MA, USA) according to providers instructions. Images were obtained with a Nikon Ri2 Color Camera and imaging with Nikon NIS Elements D Software. Six pictures (2650 x 1920 pixels) were taken at the 2, 4, 6, 8, 10, and 12 O’clock section positions.

Alcian Blue staining was analyzed with NIH Image J software for fraction (area of stained / area of the field of interest x 100%). The ages of GAG accumulation were calculated from the interception of the inoculum level with the tangent to the exponential phase as previously described [44]. The period of fast accumulation was calculated from the point at which the tangent to the exponential phase stops overlapping with the sigmoid curve.

Cross-sectional cardiomyocyte width was measured for over 50 cardiomyocytes in 15 images from three independent slides per animal stained with HE as previously reported [45] using CymoSize macro [46] and manually validated in randomly-selected images.

### Statistical analysis

To determine whether a gene or module of genes exhibited quadratic or linear association with age, we applied an extra sum of squares F test (p ≤ 0.05) to evaluate the goodness of the fit using GraphPad Prism v.8.0.2 (GraphPad Software, Irvine, CA. USA). To evaluate whether age was significant for the trait, a comparison with the line or curve (*y* = *B*_0_ + *B*_1_*x* + *B*_2_*x*^2^) with the null impact of age (*B*_1_ = 0 for linear; *B*_2_ = 0 for quadratic) and p-value ≤ 0.05 were considered significant.

Regressions for the pathways were done in a similar way using the average of the normalized and scaled expression values. For transcripts with a quadratic behavior, the vertex was also computed. The vertex of a pathway is represented as the average ± standard deviation of the vertices of all the transcripts that belong to the pathway.

The correlation matrix represents the normalized level of expression of a particular feature in a specific instance by the color of each cell, and the data were clustered by similarity with hierarchical clustering on Euclidean distances and with average linkage performed in Orange version 3.26 [47]. Row annotations include the gene and the canonical pathway. The Principal Component Analysis (PCA) was used to compute the PCA linear transformation of the input data transforming dataset with weights of individual instances or weights of principal components represented in a scatter plot.

## Results

### Morphological and blood parameters

In this study, 35 adult female baboons 7.5-22.1 years old were investigated to identify and characterize aging-related alterations in heart LV. All animals showed no age-related morphological alterations, as concluded from the absence of significant variations in body weight and body mass index (BMI), or in the abdominal, chest, waist, and hip circumferences (Figures 1B and S1). Heart weight at necropsy was consistent for all animals and independent of age (Figure 1B).

Blood biochemical parameters suggested metabolic modulation through the aging process. Both glucose (p=0.03) and triglycerides (p=0.004) circulating concentrations increased with age even though the animals were fed a healthy chow diet throughout their lifespan (Figure 1B). Total cholesterol levels in the blood show a quadratic behavior (p=0.03; Figure S1) mostly driven by the variation of LDL concentrations (p=0.03) whereas HDL concentration remained constant (Figure 1B).

### Identification of modules of genes correlated with age

Untargeted transcriptomics in the heart LV identified a total of 35,170 expressed genes that passed quality thresholds. WGCNA was used to cluster correlated genes across the samples into modules. These modules were then correlated with animal ages to determine which modules contained genes that correlated with age. Five modules (dark magenta, dark olive green, turquoise, royal blue, and dark orange) containing a total of 5,975 genes negatively correlated with age, while 6,129 genes from eight different modules (orangered 4, dark turquoise, brown, purple, orange, dark red, floral white, and salmon) are positively correlated with age (Figures 1C). Figure 1D indicates the correlation of each significative module with age, accompanied by the respective p-value and number of genes involved.

The pattern of gene expression with age is module-specific (Figure 2A). We observed that most of the modules best fit a quadratic function instead of a simple linear regression, meaning that upregulation or downregulation of the module-specific genes starts at a determined timepoint rather than occurring at a constant rate through the adult age span (Figure 2B). One exception is the purple module, which demonstrated a linear increase in gene expression with age in 7.5 to 22.1-year-old female baboons.

**Figure 2.**
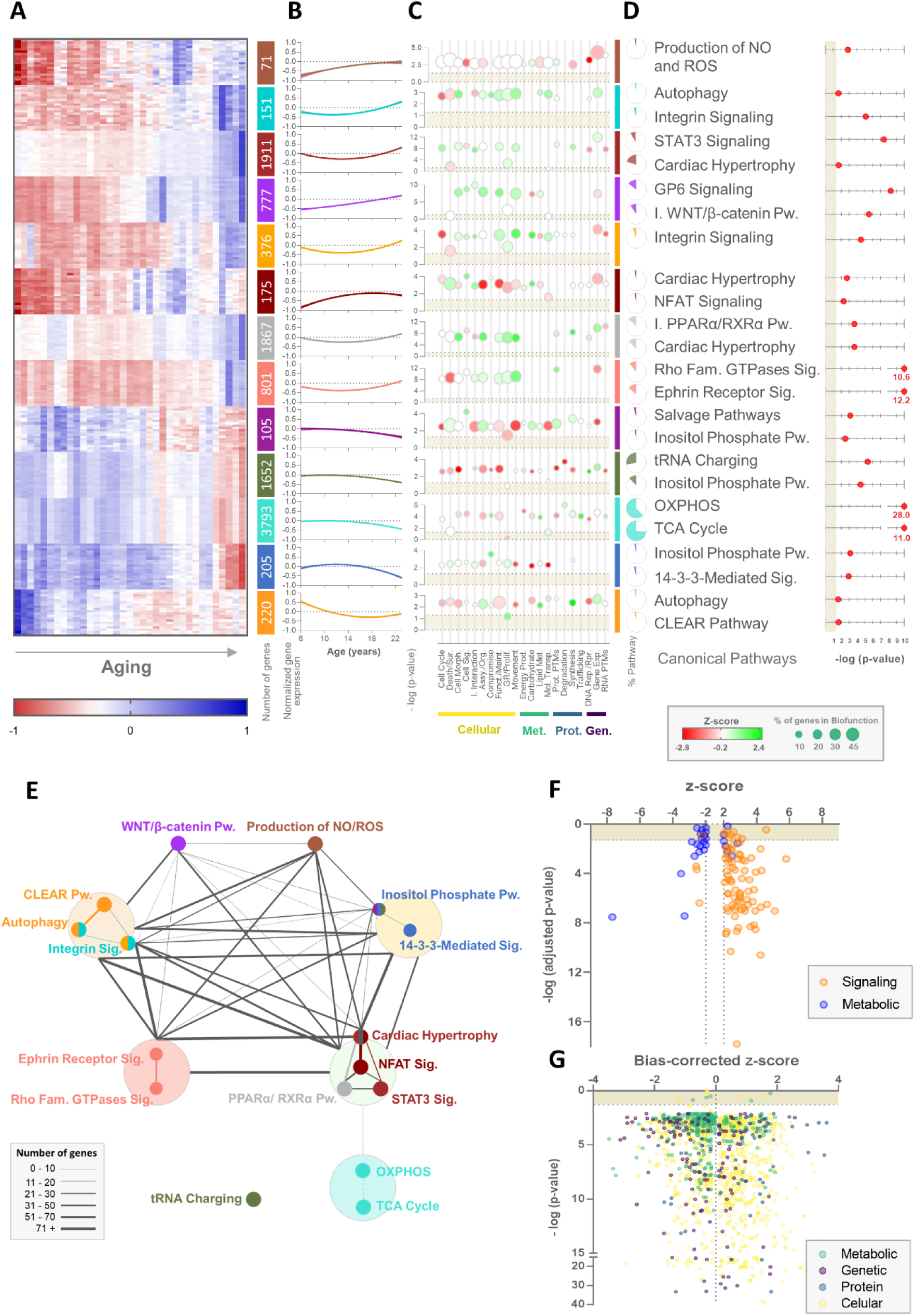
Pathway and network analysis of age-correlated modules of genes in the heart’s left ventricle. (A) Heatmap of age-related gene expression in the NHP female cardiac LV of the most significant transcripts from each module and respective number of transcripts. (B) Longitudinal profile of the average ±95% confidence interval of the gene expression of all transcripts from the module. (C) Cellular (yellow), metabolic (green), protein-related (blue), and genomic (purple) biofunctions were identified in the pathway enrichment analysis for each module. Circles represent the -log(p-value) of each biofunction for the module, their color the z-score (red negative, green positive), and the size the percentage of the genes from each biofunction identified in the respective module. (D) Most significant pathways enriched for each module, the respective -log(p-value), and the percentage of the genes from the pathway identified in the respective module. (E) The network between the pathways identified in the enrichment analysis. The color represents the color of the module, big circles contain pathways identified in the same module, and the size of the connections represents the number of genes in common between pathways. (F) Pathways identified in the module-combined pathways enrichment analysis with absolute z-score greater than 2. Dots in orange represent pathways associated with signaling functions and in blue pathways related to metabolic functions. (G) Biofunctions identified for the pathway enrichment analysis. The color of each dot represents cellular (in yellow), metabolic (in green), protein-related (in blue), and genomic (in purple) biofunctions. Assy. – assembly; Exp. – expression; Fam. – family; Funct. – function; GR – growth; I. – inhibition; Maint. – maintenance; Met. – metabolism; Mol. – molecular; Morph. – morphology; Org. – organization; OXPHOS – oxidative phosphorylation; Prod. – production; Prolif. – proliferation; Prot. PTMs – protein post-translational modifications; Pw. – pathway; Rep. – replication; RNA PTMs – RNA post-transcriptional modifications; Rpr. – repair; Sig. – signaling; Sur. – survival; TCA - tricarboxylic acid; Transp. – transport.

### Pathway and network analysis of the modules of genes correlated with age

To better understand how variation in gene expression in the age-associated modules may affect LV heart cellular function, we performed pathway enrichment analysis for every significantly age-correlated module. The pathways and biological functions associated with each module partially overlap but each has specific signatures (Figure 2C). Cellular functions are mostly upregulated in modules positively correlated with age. In fact, the most significant (*i.e.*, p-value and z-score) canonical pathways identified in these modules (Figure 2D) are related to alterations in cellular morphology (Integrin Signaling–dark turquoise and orange modules; Rho Family GTPases Signaling–salmon module; GP6 (Glycoprotein VI) Signaling–purple module; Ephrin Receptor Signaling–salmon module), cellular growth (Cardiac Hypertrophy– brown, dark red, and floral white modules; STAT3 (signal transducer and activator of transcription 3) Signaling–brown module; NFAT Signaling–dark red module), and heart development and homeostasis (inactivation WNT/β-catenin–purple module).

Expression of transcripts and pathways related to metabolic functions are generally in modules downregulated with age and most of the metabolism-related genes are found in modules that are negatively correlated with age. Metabolic canonical pathways are the most significant among these modules with considerable overlap of oxidative phosphorylation (OXPHOS) (55.9%), TCA Cycle (73.9%), and the Inositol Phosphate Pathway (more significant in dark magenta, dark olive green, and royal blue modules). Some modest positive z-scores for other pathways related to metabolic functions are also observed in the dark turquoise and orange modules, even though they are not represented in the most significant canonical pathways. Altogether, gene co-expression related to metabolic functions appears to decrease with age impacting multiple pathways vital for cardiac energy production.

Interestingly, most of these canonical pathways overlap and are mostly associated with cardiomyocyte structural alterations and cardiac remodeling (Figure 2E). Indeed, when we combined the gene lists from modules with the same directionality (upregulated or downregulated with age) and did a subsequent pathway enrichment analysis, we identified the same pathways. In total, 108 canonical pathways were identified with z-scores greater than 2 or lower than −2 (Figure S2A). The canonical pathways are classified into two categories: metabolic and signaling pathways. By combining the modules it is possible to obtain a better overview of the heart’s LV transcriptional modulation of cellular and molecular functions throughout aging. Once again, the identified canonical pathways downregulated with aging are primarily related to metabolism (20 out of 24, Binomial test: p = 0.002; Figure 2F). Consistent with these findings, only 7 in 84 of the upregulated canonical pathways are related to metabolism (Binomial test: p < 0.0001).

Analysis of the biological functions associated with gene expression that correlated with age provides a similar perspective. The most significantly upregulated functions associated with age include cellular development, movement, cell-mediated immune response, protein post-translational modifications, and protein trafficking (Figure S2B). As previously observed, the majority of biological functions related to cellular maintenance and processes are upregulated (present significant positive z-scores) with age. On the other hand, the downregulated biological functions include mostly protein- and metabolic-related functions, namely DNA replication and repair, RNA post-transcriptional modifications, small molecule biochemistry, and lipid and nucleic acid metabolism. This is in line with the pathway enrichment analysis where the majority of canonical pathways identified as negatively correlated with age belong to metabolic pathways and functions. Indeed, while cellular, genetic, and protein functions are nearly equally distributed according to the biased-corrected z-score, the metabolic-related biological functions clearly cluster on the negative side of the axis (Binomial test: p < 0.0001; Figure 2G). This again suggests a consistent downregulation of the pathways related to metabolism on the cardiac LV of female baboons with age.

Two major cellular cardiac alterations stand out: a general downregulation of metabolic pathways and an upregulation of signaling pathways related to cardiac remodeling. Taking this into consideration, we analyzed each metabolic pathway individually to understand whether and potentially how the modulation of cellular metabolism in the LV is related to cardiac remodeling signaling pathways.

### Cardiac metabolic adaptations with aging

Cardiomyocyte metabolism and energy production are primarily dependent on FA oxidation, a process that includes FA uptake, activation and β-oxidation, and subsequent metabolism of intermediates through the TCA cycle and OXPHOS [48]. At least half of the transcripts from genes belonging to these pathways were downregulated (OXPHOS: 62.6% transcripts of 62 genes; TCA cycle: 71.4% transcripts of 16 genes; FA β-oxidation: 59.3% transcripts of 16 genes; FA uptake/activation: 50% transcripts of 9 genes) with age (Figure 3A).

**Figure 3.**
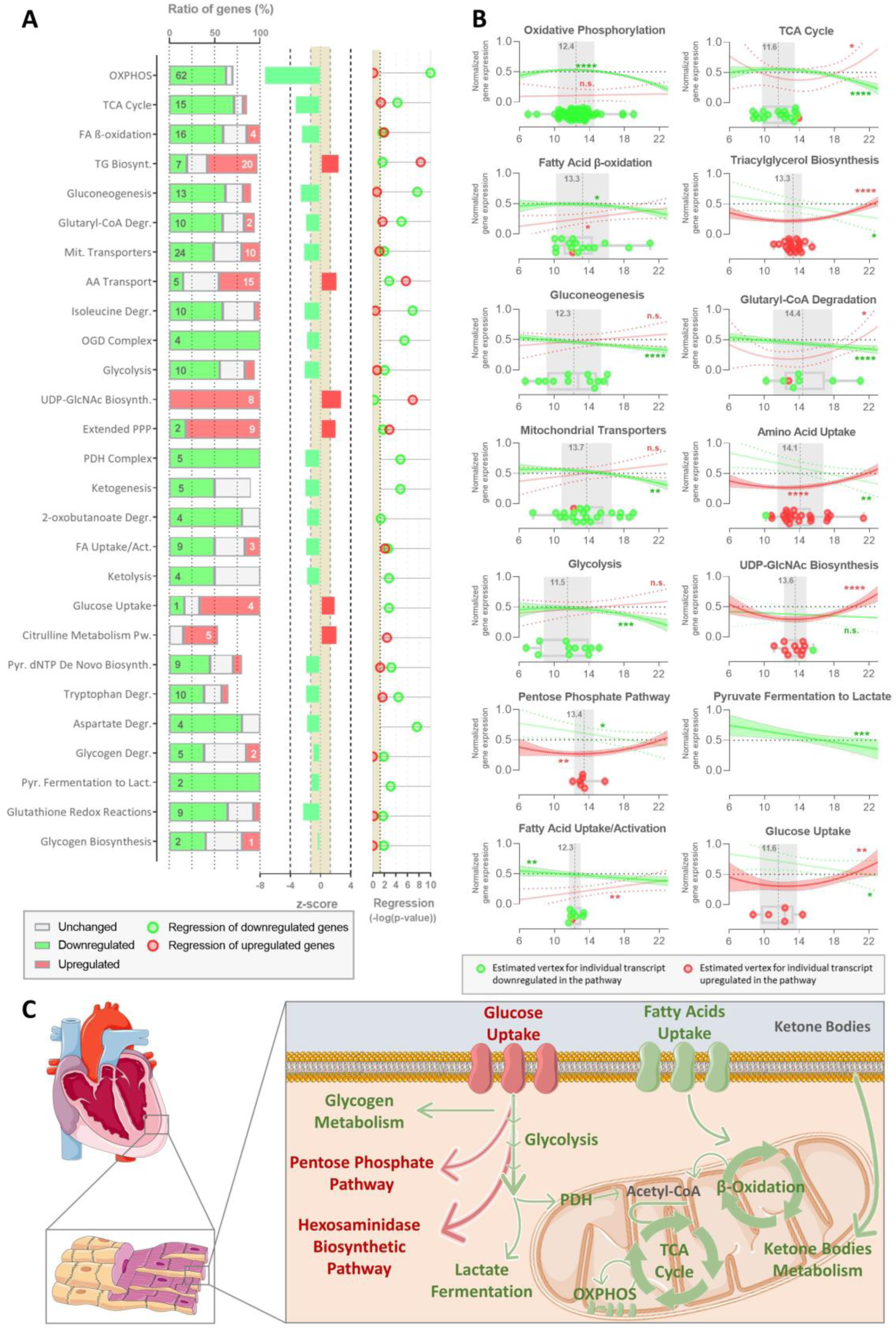
Pathway enrichment analysis of the transcripts that correlate with age and are involved in the NHP female cardiac metabolism. (A) The pathways related to metabolism identified in the pathway enrichment analysis are depicted. The ratio of the genes from each canonical pathway identified and directionality (downregulated – green; upregulated – red; unchanged – grey), Z-score from the respective pathway, and the -log(p-value) of the regression of the relative transcripts expression with age for the downregulated and upregulated genes. (B) Longitudinal profile of relative transcript expression with age, boxplot with transcripts tipping points (red – upregulated, green – downregulated), and the average age of the tipping point (grey) from the metabolic canonical pathway. (C) Schematic representation of the relation between the identified pathways. Downregulated metabolic pathways are represented in green and upregulated mechanisms in red. AA – amino acid; Act. – activation; Biosynth. – biosynthesis; Degr. – degradation; Lact. – lactate; Mit. – mitochondrial; OXPHOS – oxidative phosphorylation; PDH – pyruvate dehydrogenase; PPP – pentose phosphate pathway; Pw. – pathway; Pyr. – pyrimidine; TCA - tricarboxylic acid.

Even though lower FA utilization to produce energy in the heart is commonly compensated by the rise of oxidation of other substrates, glucose-(glycolysis: 55.6% transcripts of 10 genes), ketone bodies-(ketolysis: 50% transcripts of 4 genes), and amino acid-related (isoleucine, tryptophan, and aspartate degradation) oxidation pathways are also downregulated with age in heart LV of female baboons.

Although glucose utilization is reduced, the expression of glucose transporter genes is upregulated except for the insulin-dependent transporter SLC2A4 (Figure 3). Yet, transcripts associated with glucose conversion to glycogen storage are also downregulated. Interestingly, the expression of genes of two metabolic pathways channeling glucose utilization is upregulated with age: pentose phosphate pathway (PPP: 81.8% transcripts of 9 genes) and hexosamine biosynthetic pathway (HBP; *i.e.* UDP-GlcNAc Biosynthesis: 100% transcripts of 8 genes). Other upregulated canonical pathways include the anabolic synthesis of triglycerides (55.6% transcripts of 20 genes) and the citrulline pathway (38.5% transcripts of 5 genes).

Surprisingly, gene expression associated with these canonical pathways is not always linear but rather better fits a quadratic function (Figure 3B). This relationship was observed in downregulated pathways for, but not exclusively, OXPHOS, TCA cycle, FA β-oxidation, and glycolysis. Accordingly, the upregulation of glucose uptake, PPP, and HBP also follows a quadratic regression. All of these pathways present a constant gene expression with age until female NHP reached 12.93±0.32 years old when these canonical pathways change the trajectory and start decreasing or at 13.32±0.36 years old when gene expression starts increasing.

Overall, our results indicate that transcripts relevant to cardiac metabolism are modulated during the normal (healthy) aging process resulting in a general decrease in the catabolic pathways and an increase of anabolic pathways at specific ages mostly related to glucose intermediate metabolism (Figure 3C).

### Glycosaminoglycans accumulation

Stimulation of the UDP-GlcNAc biosynthesis via HBP can result in increased O-GlcNAcylation and the synthesis of GAGs. Glycosaminoglycans are a family of polysaccharides that are distributed at the cell surface or in the ECM attached to core proteins (PGs), or released while synthesized as in the case of the hyaluronan [25]. Of the four different types of GAGs, it is important to highlight that Chondroitin/Dermatan biosynthesis is one of the metabolic canonical pathways correlated with age in our analysis (Figure S2A). To examine if the cardiac metabolic shift is associated with increased GAG synthesis, we focused on the expression of genes associated with GAG-related precursor synthesis, intermediate metabolism, and GAG synthesis, modification, and degradation. We performed cluster analysis of the genes previously identified as upregulated or downregulated with age (Figure 4A). Interestingly, this analysis clustered only genes associated with GAG degradation. The other cluster included the genes of all the other classes, suggesting a similar age-related longitudinal profile among them. These findings were corroborated by a Principal Component Analysis of the same genes (Figure 4B).

**Figure 4.**
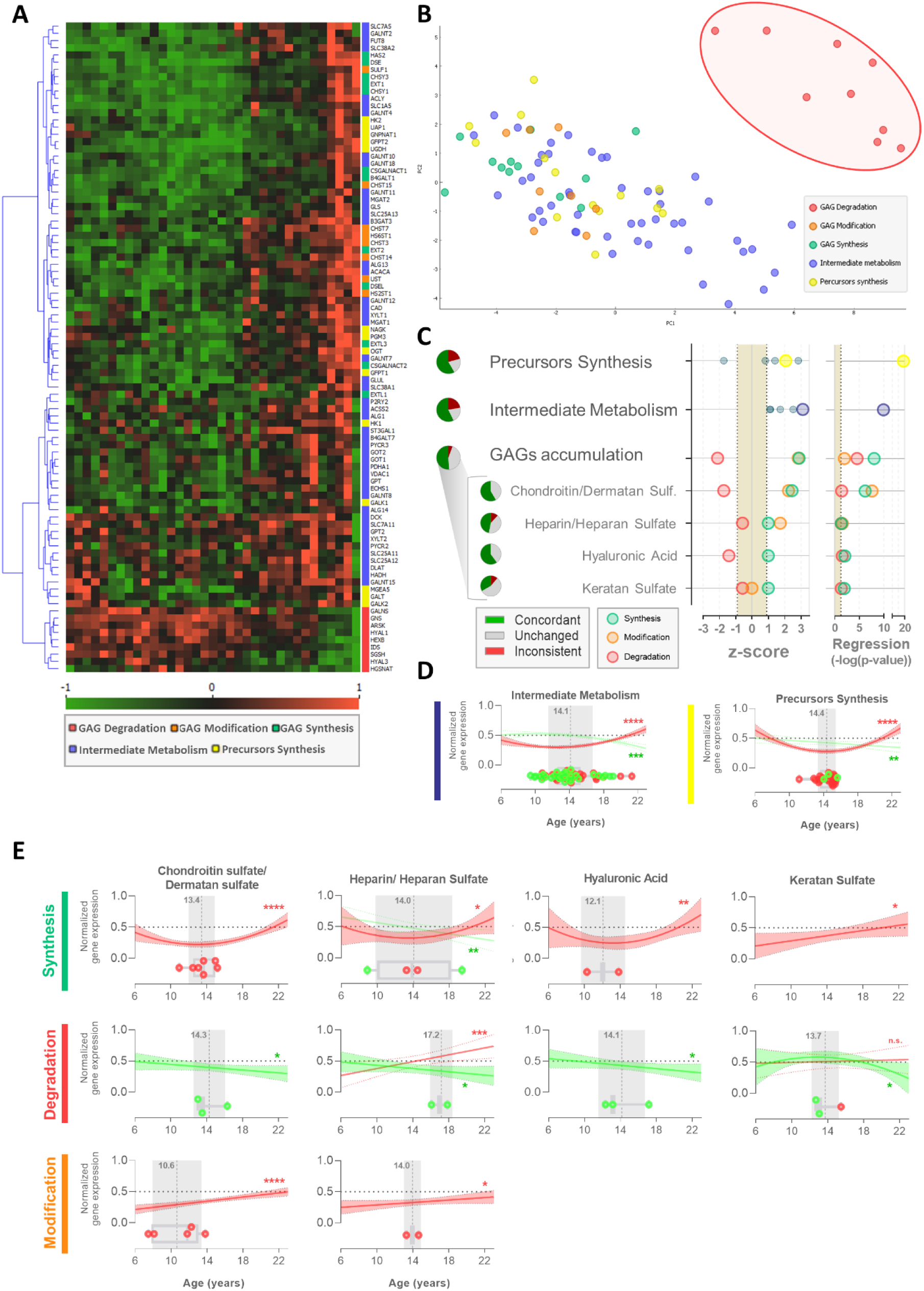
Glycosaminoglycan-related pathways transcriptomics in heart with aging. (A) Heatmap showing the longitudinal relative gene expression of transcripts related to GAG degradation (red), GAG modification (orange), GAG synthesis (green), intermediate metabolism (blue), and precursors synthesis (yellow). (B) Principal component analysis of the age-associated profile of gene expression. (C) Z-score of the glycosaminoglycan-related pathways from their transcript expression, percentage of the genes from the pathway with a concordant (green), inconsistent (red) and unchanged (grey) behavior with age, and circles represent the pathway’s -log(p-value) of regression. Transcripts associated with GAG synthesis in green circles, with GAG modification in orange circles, and with GAG degradation in red circles. (D) Longitudinal profile of the variation of the normalized transcript expression with age for the proteins associated with intermediate metabolism and precursor synthesis, and (E) with the synthesis, degradation, and modification of each type of glycosaminoglycan. Boxplots with transcripts tipping points, in red, represent upregulated transcripts, in green downregulated transcripts, and the age of the average tipping point in grey. GAG – glycosaminoglycan; Sulf. – sulfate.

GAG-related precursors synthesis includes genes in the metabolic pathways associated with UDP-GlcNAc, uridine diphosphate N-acetylgalactosamine (UDP-GalNAc), D-glucuronic acid (GlcA), and Galactose synthesis. These genes present an overall positive z-score (Figure 4C) and a significant increase in their expression with age (p<0.01) after 14.21±0.30 years old in female baboons (Figure 4D). Nonetheless, the genes related to galactose synthesis (required for KS synthesis) present a negative z-score while all the other precursors show a positive z-score. Genes related to intermediate metabolism (i.e., glutamate/glutamine, acetyl-CoA; UTP, and protein backbone preparation to glycosylation) present increased expression with age (p<0.01) after 14.29±0.37 years old in female baboons and positive z-score, suggesting that the heart LV metabolism modulation is promoting these metabolites pools and, consequently, contributing to GAGs synthesis.

The general overview of the genes related to GAG metabolism shows an increase in the expression of transcripts associated with GAG synthesis (z-score=2.9; p<0.01) and modification (*i.e.*, sulfation; z-score=2.8; p<0.01) with age, while genes related to GAG degradation show decreased expression with age (z-score=-2.1; p<0.01). Altogether, cardiac gene expression appears to favor increases in GAG numbers or the length of the polysaccharide chains. This is consistent for the four classes of GAGs (Figure 4E), except for gene expression related to KS modification which showed no alteration in expression throughout aging. Indeed, genes specifically related to KS present modest z-scores (synthesis: z-score=1; degradation: z-score=-0.6; modification: z-score=0). Together with the downregulated metabolism of galactose, data suggest that KS is the GAG least impacted by age in the female heart’s LV.

Metabolism related to CS/DS and HA presents the most consistent z-scores for each function, more than 50% of genes are upregulated in the GAG pool, with no contradictory gene expression (Figure 4C). Even though almost half of the genes are concordant with HS accumulation, one gene related to HS synthesis is downregulated and another responsible for HS degradation is upregulated. Nonetheless, overall the transcript profiles are consistent with increased synthesis of CS/DS (p<0.01), HA (p<0.01), and HS (p=0.04) with age. The quadratic analysis suggests that the cardiac alteration of the gene expression related to GAG metabolisms starts at 13.65±0.57 (Figure 4E). Transcripts associated with GAG degradation of CS/DS (p=0.04), HA (p=0.03), and HS (p=0.04) and for CS/DS (p<0.01) and HS (p=0.03) sulfation appear to be decreasing with age. It is important to note that transcript expression of PG is also upregulated for decorin, betaglycan, versican, and lumican with age, but remains unaltered for the other PGs.

Alterations in gene expression related to GAG metabolism suggest GAG accumulation with age in the ECM of the cardiac LV. This was indeed confirmed with Alcian blue staining histological analysis (Figure 5A). Age-related GAG staining follows a sigmoidal curve (Figure 5B), which allows the estimation of two phases of accumulation: the total range of accumulation between 11.3 to 21.3 years old (∼45–85 human equivalent years), and the interval of faster accumulation between 13.5 to 18.0 years old (∼54–72 human equivalent years). GAG accumulation is generally associated with increased ECM stiffness, which is commonly seen with cardiac remodeling, ECM-induced intracellular signaling, and cardiac hypertrophy [49,50].

**Figure 5.**
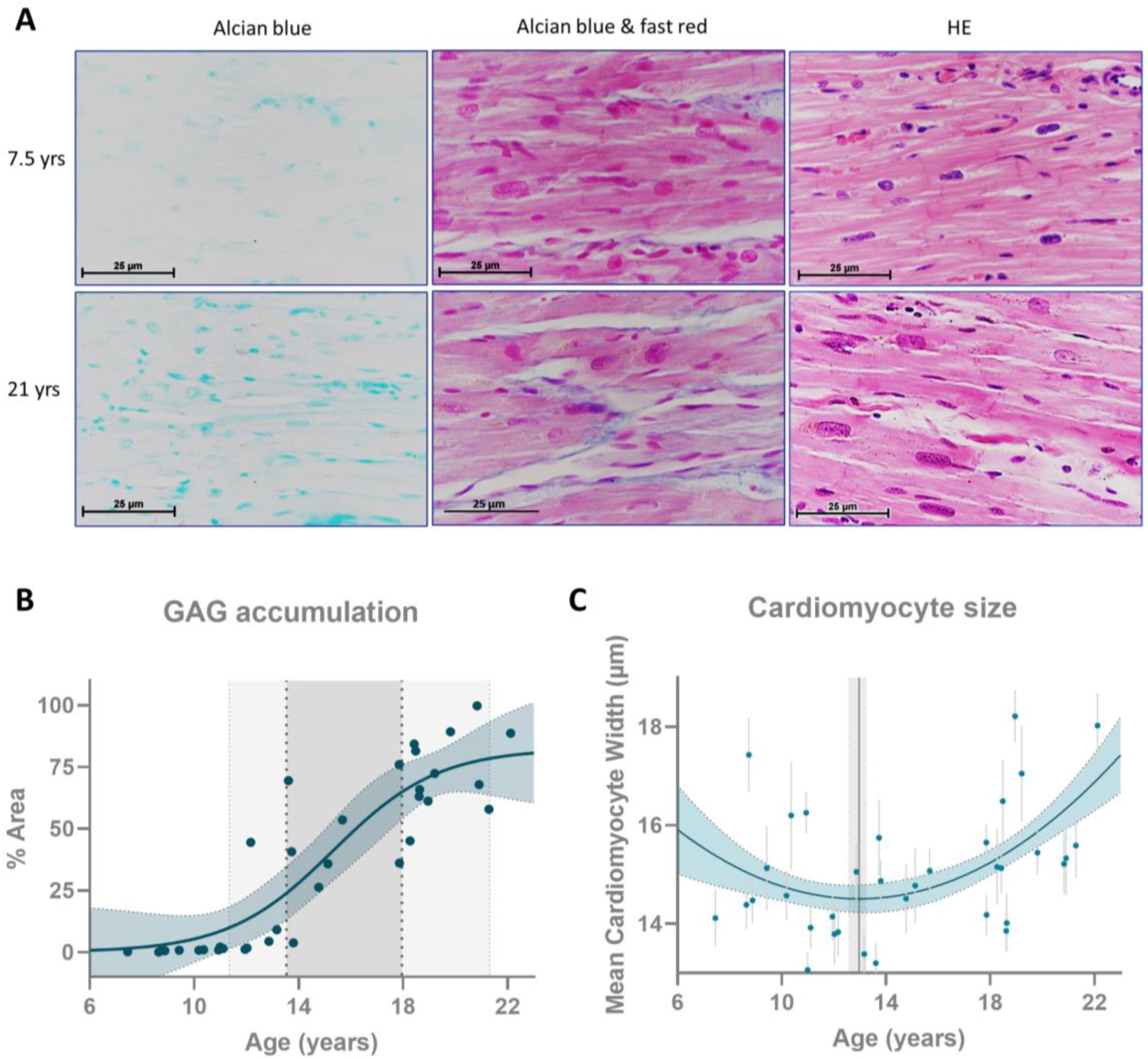
Glycosaminoglycan accumulation with age in the NHP female heart’s left ventricle. (A) Histological staining of glycosaminoglycans and hematoxylin and eosin in heart tissue, (B) respective quantification of glycosaminoglycan with age (depicted ages of GAG accumulation in light grey and of fast accumulation in dark grey), and (C) variation of left ventricle cardiomyocyte size with age (each dot represents the mean±95% confidence interval for each individual; grey line the age of the vertice calculated from the quadratic curve). GAG – glycosaminoglycan; HE - hematoxylin and eosin staining.

### Age-related cardiac remodeling

The development of cardiac hypertrophy is considered a hallmark of aging [16]. Nevertheless, whether this phenotype is a trigger for age-related CVD or merely a consequence of a cascade of events is not clear. We measured cardiomyocytes’ cross-sectional width using hematoxylin-eosin stain to assess cardiac hypertrophy. We identified an increase in cardiomyocyte width, which was first evident at 13.0 years old (∼52 human equivalent years), 95% CI [12.6, 13.2]. This increase in cardiomyocyte width followed a quadratic growth pattern. The development of cardiac hypertrophy with age is consistent with the canonical pathway identified in our transcriptomics analysis. The pathway-enrichment analysis identified seven signaling pathways related to cardiomyocyte growth with z-scores greater than two (Figure 6A). The vertices of the transcripts associated with these pathways show that the change in gene expression trajectory occurs at 14.63±0.18 years in female baboons (∼58 human equivalent years) for the pathways related to the development of cardiac hypertrophy with age.

**Figure 6.**
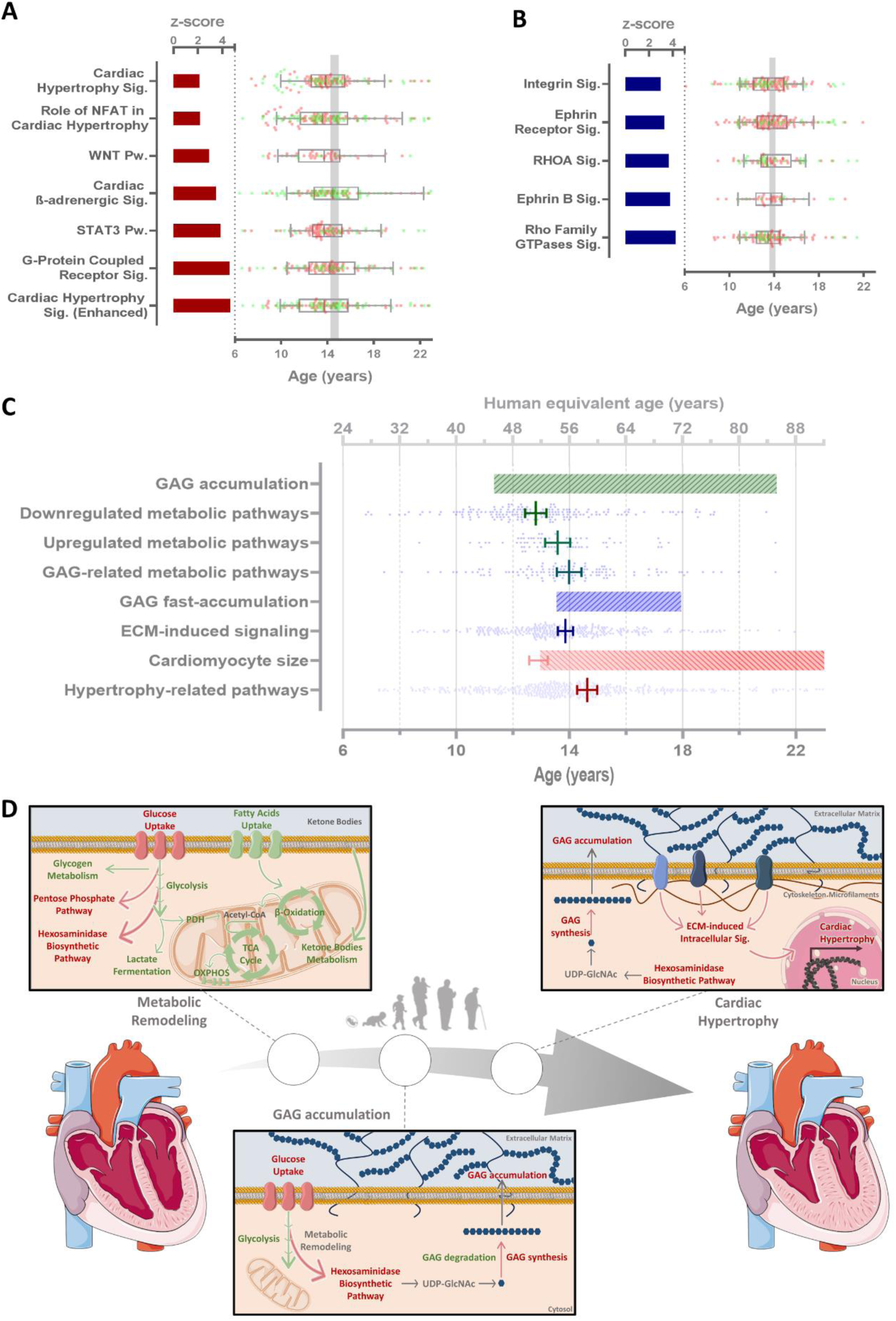
Temporal progression of age-related molecular events in the NHP female heart’s left ventricle. (A) Cardiac hypertrophy-related and (B) ECM-induced pathways with absolute z-score greater than 2. Bars represent the z-score and boxplot the vertices of the transcripts with quadratic behavior from each pathway (in red transcripts upregulated with age and in green transcripts downregulated with age). (C) Comparison of baboon ages (and human equivalent age) from the calculation of the tipping points predicting the ages at which the transcriptomics aging-related changes for different pathways, periods of GAG accumulation, fast accumulation, and increased cardiomyocyte size. (D) Schematic representation of the novel molecular proposed mechanisms associated with cardiac aging. ECM – extracellular matrix; GAG – glycosaminoglycan; OXPHOS – oxidative phosphorylation; PDH – pyruvate dehydrogenase; Pw. – pathway; Sig. – signaling; TCA - tricarboxylic acid.

To evaluate the relationship between GAG accumulation and cardiac remodeling, we evaluated how the signaling pathways related to ECM modulation change with age. Generally, ECM-induced signaling increases with age (Figure 6B) and most of the genes fit a quadratic curve in which the gene expression starts increasing at ∼14 years (approximately 56 human years). When compared to other cellular mechanisms associated with age, upregulation of the ECM-induced signaling occurs at a similar age when GAGs begin rapidly accumulating in the ECM, matching the vertices of transcripts from GAG-related metabolic pathways (Figure 6C). These alterations appear to follow the downregulation of cardiac metabolism and precede the upregulation of genes in hypertrophy-related pathways.

Collectively, our results demonstrate a strong correlation between gene expression in female primate heart LV and age (34.4% of the transcripts). Transcript profiles related to cardiac metabolism, mostly catabolic pathways like FA oxidation, glycolysis, and ketone bodies metabolism, were largely consistent with metabolic downregulated as one of the earliest observable changes during aging. Nonetheless, the expression of some genes related to metabolic pathways increases with age, including HBP and PPP, critical to GAG synthesis. The most evident GAG metabolism-associated gene expression alterations appear to favor increased numbers and/or length of CS/DS and HA (potentially also HS) chains by upregulating synthesis pathways and downregulating degradation pathways, thus promoting GAG accumulation. These cardiac adaptations are coincident with the ECM-induced pathways and precede cardiac hypertrophy-related gene expression (Figure 6D).

## Discussion

Worldwide demography has changed over recent decades with increased median age and proportion of older adults [51]. Aging has been considered one of the major risk factors for CVD, and the recent demographic trend accentuates the overall number of individuals living with CVD [52]. Other risk factors for CVD are circulating triglycerides, total cholesterol, LDL and HDL cholesterol [53]. In our study, in which 35 female baboons were studied across the adult age span, we showed that glucose, triglyceride, and total and LDL cholesterol circulating concentrations were significantly associated with age in opposition to HDL cholesterol. The linear increase in glucose and triglycerides is consistent with previous reports in women [54,55]. In baboons, LDL blood concentrations vary in a quadratic profile with higher levels at middle-adult ages. Our study of animals living in a controlled environment fed a healthy diet is consistent with lipoprotein and morphological changes seen with aging in women [56].

Cardiac aging in NHP recapitulates physiological, cellular, and molecular mechanisms of normative aging in humans [57]. Due to the inability to collect cardiac LV tissue from healthy humans, cardiac studies in humans are frequently conducted on non-healthy individuals without CVD-related disorders or in cases of sudden death and accident victims. Nevertheless, tissue collection in such cases may interfere with the quality of the analysis. It is worth noting, that we attempted similar analyses on human heart transcriptomic data available at the Genotype-Tissue Expression Portal. However, our bioinformatic approach did not reveal any age-correlated transcripts, potentially due to variability resulting from disparity in post-mortem intervals, life style, and other contributors. Thus, the use of NHP represents a unique opportunity to study molecular signatures of physiologic heart aging in healthy primates with relevance to humans. We performed a comprehensive statistical and pathway-enrichment analysis on untargeted transcriptomes in LV heart from a longitudinal age-related female baboon cohort.

Cardiomyocyte aging includes multiple cellular adaptations to the age-related accumulation of defective molecules, protein dysfunction, and impairment in reparative mechanisms [58,59]. Cardiac remodeling is a physiological process that is initially compensatory, but when sustained can be detrimental to cardiac function [3]. Age-associated increased cardiac LV wall thickness is a consequence of the elevation in afterload or the pressure against which the heart must work to eject blood [16]. The progressive loss of cardiomyocytes and hypertrophy of those remaining increases inflammation and cardiac fibrosis while compromising heart pumping capacity and interfering with cardiomyocyte electrical coupling [58]. We previously observed age-related decreased LV ejection fraction and rate, loss of myocardial mass, and augmented LV end-diastolic sphericity index in female baboons [18,19]. Consistent with the imaging results, we found increased cardiomyocyte width in addition to upregulation of cardiac hypertrophy-related gene expression and associated canonical pathways including STAT3 and NFAT signaling. The upregulation of STAT3 transcriptional activity via SIRT2 downregulation was previously reported in geriatric NHP along with increased cardiomyocyte enlargement and sarcomere structural abnormalities [60]. Interestingly, the age at which cardiomyocyte width begins to increase precedes the inflection points of gene expression associated with cardiac hypertrophy. It is possible that altered expression of those genes regulates maladaptive cardiac hypertrophy resulting from GAG accumulation and that the initial increase of cardiomyocyte size results from a natural compensatory mechanism concomitant with the downregulation of metabolism-related genes.

Due to the elevated demand for oxygen and energy on hypertrophic cardiomyocytes, heart remodeling includes modulation of the ECM, which can regulate the extracellular milieu and other systemic factors (*e.g.*, inflammation, growth factors) [58,61]. Dysregulated ECM homeostasis in aged hearts leads to cardiac fibrosis [16] in multiple species including mice [62], rats [63], dogs [64], and humans [65]. As compensation for cardiomyocyte loss, the ECM content increases in a process mostly controlled by the balance of ECM constituent synthesis and degradation [21,58]. We showed here for the first time that the machinery of synthesis of GAGs, related intermediate metabolism, and availability of precursors are upregulated transcriptionally in a process temporally coincident with GAG increased accumulation rate with normative aging in the NHP female cardiac LV. Since GAG regulation is delicately controlled by synthesis and degradation enzymes, and performed by the different myocardial cell types [21], our findings in heart bulk transcriptomics unveil hints of transcriptionally regulated age-related GAG accumulation. Indeed, overall gene expression of GAG degradation enzymes is decreased with age. Previous reports show higher LV total GAG concentrations in failing pediatric and adult hearts, chronic stress, and pressure-overload cardiac remodeling [21,22,26,66,67]. Nevertheless, our study demonstrates that GAG accumulation is transcriptionally regulated and occurs as a natural consequence of physiological aging. Additionally, previous studies have indicated that GAGs tend to accumulate in disease-prone areas of the vascular system, such as branch points, aligning with lipid deposition [67]. Consistent with these findings, our earlier research reported an increase in apical pericardial fat in aging female baboons [17].

Higher CS and HS levels are aligned with increased LV posterior wall thickness and enlarged average cardiomyocyte size due to chronic stress in mice [22], similar to our findings. Indeed, CS accumulates in the LV end-stage cardiomyopathic heart in humans and in a rat model of pressure-overload cardiac remodeling [21]. The existing literature consistently suggests that GAG accumulation is involved in heart pathophysiology. Nevertheless, our novel findings indicate that GAG accumulation may occur prior to cardiac hypertrophy, play a role in physiological aging, and may directly contribute to the development of CVD.

Congenital heart and vascular defects (*e.g.*, heart septal defect, bicuspid aortic valve, and aortic root dilatation) are found in individuals with mutations in enzymes related to GAG metabolism [25,68–72]. Genetic variance in GAG-related genes appears to be linked with multiple cardiovascular dysfunctions (Table 1) [73–97]. Proteoglycans rich in GAG were found to be increased in the heart’s ECM in CVDs [98–101]. Indeed, increased versican protein was reported, when comparing pediatric with adult control hearts [26], and lumican knock-out mice show higher susceptibility to aging and increased mortality due to reduced systolic function during the age-induced structural remodeling of the heart [102]. GAG sulfation patterns critically affect the binding affinity to other proteins such as anti-thrombin III, heparin cofactor 2, bFGF, heparin-binding protein, and transforming growth factor-β (TGF-β) [22,26,27,103–106]. GAG sulfation and numbers can directly modulate inflammation-related processes [107]. Corroborating our findings, an age-dependent increase of GAG sulfation in the LV, but not in circulation was previously described [66].

**Table 1.**
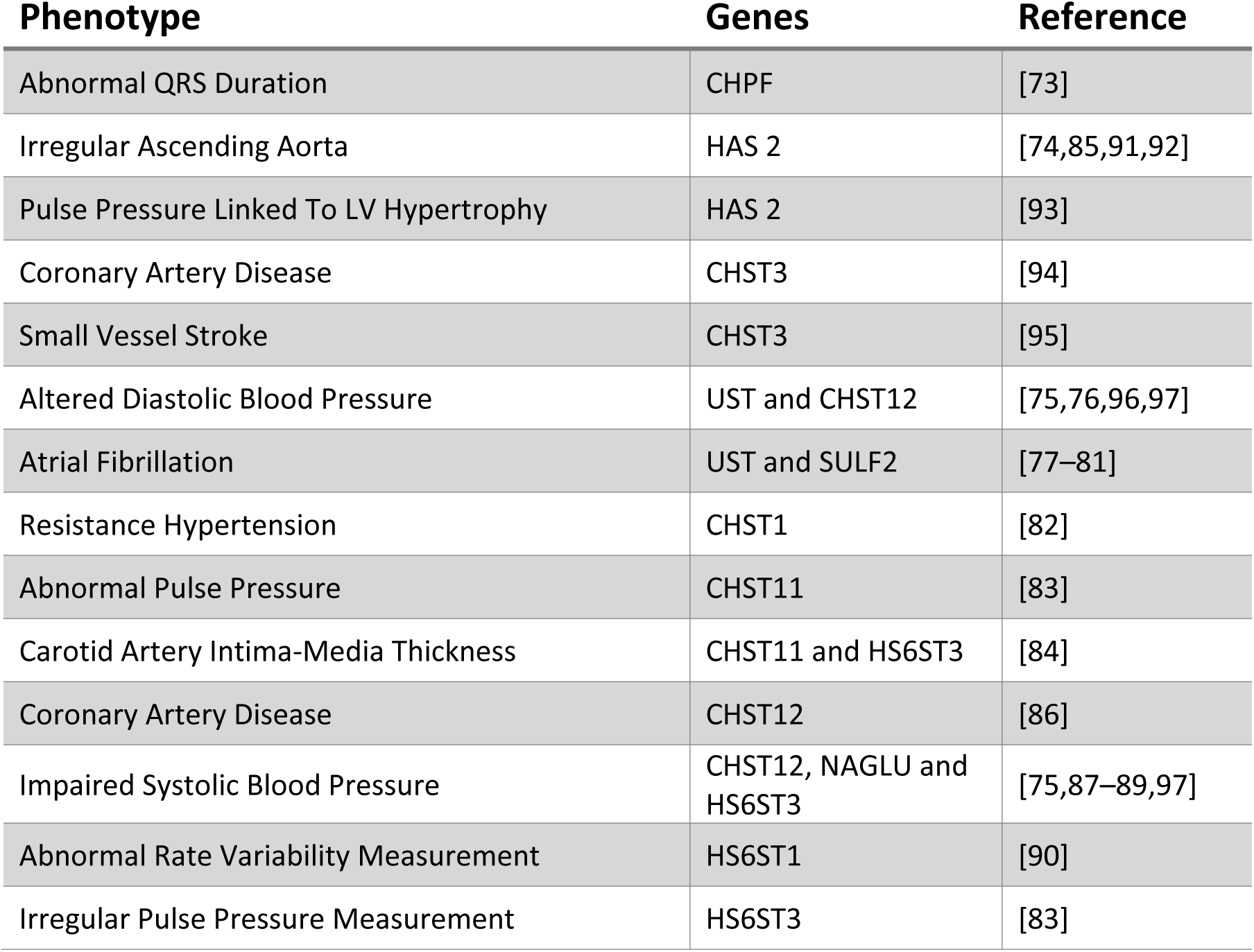
Cardiovascular phenotypes associated with genetic variance in glycosaminoglycan metabolism, synthesis, and degradation.

Besides their role as an extracellular reservoir, GAGs also modulate tissue stiffness. Hyaluronan is of major importance for cardiac ECM remodeling and the mechanical integrity of the matrix [107]. In this study, we found transcriptional upregulation of the enzymes responsible for HA synthesis and downregulation of degrading enzymes, suggesting HA accumulation. In addition, we suggest a cardiac metabolic shift that may promote glucose consumption through HBP. The final product of HBP is UDP-GlcNAc whose intracellular concentration directly regulates the rate of HA synthesis [108]. Deficiency in hyaluronidase 2 is linked to heart failure [109] and HAS2 deletion causes embryonic lethality due to cardiac malformation [110]. While HA-binding proteins can modulate ECM’s pro-inflammatory state, and recruitment and retention of other cell types (e.g., fibroblasts, macrophages) [111], increased HA accumulation is associated with increased ECM stiffness [49] while syndecan-4 knockout mice display reduced myocardial stiffness [112]. These observations reinforce our previous findings, which indicated a decline in the LV cardiac index, normalized myocardial mass, normalized average filling rate, and ejection fraction, along with cardiac remodeling due to a higher LV end-diastolic sphericity index in aging female baboons [18–20].

It was previously proposed that GAG number, composition, and sulfation in the heart allow the ECM to serve as an extracellular reservoir for growth factors, hormones, and cytokines [22,26,103–105]. Our results document GAG accumulation in the heart’s ECM as well as an upregulation of gene expression of the sulfation machinery, which increases the negative charge of the proteoglycans and modulates the cardiac cellular microenvironment and consequent intercellular communication and other external signaling pathways [24]. In addition, we found evidence for increases in some PG core protein expression (*i.e.*, decorin, betaglycan, versican, and lumican) but not all PGs, which are required for the synthesis of DS, CS, HS, and KS. It is possible that more than PG levels, increases in overall numbers and or lengths of GAG chains are occurring with age. Concomitant with GAG accumulation, we report increased transcript levels for ECM-induced signaling pathways, suggesting a role in cellular function regulation from GAG accumulation.

Heart ECM stiffness can transfer physical signals to the intracellular matrix through mechanical conduction, modulating intracellular signaling pathways [49]. Among the key players in the relationship between ECM stiffness and intracellular signaling, Integrin and Rho GTPases signaling emerge as two major components [49,103]. These two canonical pathways were identified as upregulated coincident with the period of GAG fast accumulation, corroborating the hypothesis of an age-associated increase in cardiac ECM stiffness due to GAG accumulation and modification. Cardiomyocytes and endomysial fibroblasts sense the ECM network through integrin ECM receptors, which are activated by the engagement of ECM ligands regulating intracellular signaling. Downstream effectors include PI3K/Akt signaling via FAK and the promotion of GLUT1 and GLUT3 gene expression [103,113,114] or the increased expression of amino acid transporters [49,115]. In concordance, we show upregulated transcripts for amino acid transporters and GLUT1 and GLUT3 (but not GLUT4) with aging in the heart LV of female baboons. Through Akt, integrin receptors can also regulate lipid metabolism by promoting triglyceride biosynthesis [116], which we also identified as a canonical pathway stimulated with aging in the female heart’s LV. ECM stiffness can also impact cellular metabolism through Rho/ROCK signaling by affecting the actin cytoskeleton that regulates glycolysis-enzymes activities [117–119].

Indeed, multiple studies report cardiac metabolism adaptations along with aging namely resulting in the loss of metabolic flexibility, mitochondrial efficiency, and ATP depletion in rodents [6,9–12,48]. We observed lower expression of mitochondrial-related genes for OXPHOS and FA oxidation with aging as previously reported [10,11] including in NHP [60]. As a compensatory mechanism, previous studies suggest a metabolic shift that favors glucose oxidation instead of FA oxidation resembling fetal cardiomyocyte-like metabolism, in a process transcriptionally regulated [11,12,120]. In this study, we observed an increase in glucose transporters gene expression however, genes related to glucose oxidation (e.g., glycolysis) and storage (e.g., glycogen biosynthesis and degradation) are downregulated. In fact, in our analysis, almost all the canonical pathways downregulated with age are related to metabolism and those are the earliest alterations detected in the cardiac LV tissue with age preceding GAG accumulation and cardiac hypertrophy. This suggests that cardiac metabolism slows throughout aging, compromising heart pumping, and might be the cause of decreased ejection fraction and rate in these animals [18]. It was proposed that lower FA uptake and oxidation and an increase in glucose utilization accelerate the development of cardiac hypertrophy, yet the underlying mechanisms are not well understood [8,14]. We observed higher expression of genes related to the pentose phosphate pathway and HBP which channel glucose consumption that may promote GAG accumulation and later cardiac hypertrophy.

HBP flux is mostly dependent on nutrient availability through the uptake and intermediary metabolism (glutamine, acetyl-CoA, UTP) [121]. While acute HBP activation increases UDP-GlcNAc and O-GlcNAcylation levels and FA oxidation [122], HBP chronic upregulation under hemodynamic stress induces pathological cardiac hypertrophy and heart failure [13], similar to our results. The final product of HBP, UDP-GlcNAc, can undergo the synthesis of GAG or be used for O-GlcNAcylation [121]. Moreover, persistent induction of GFPT1, the first and rate-limiting enzyme of HBP, in the heart can activate mTOR signaling whose role in both physiological and pathological hypertrophic growth is well-described [50]. Indeed, cardiac hypertrophic growth generally requires the activation of multiple signaling pathways including PI3K/AKT, mTOR, and p70S6K [30] which were all identified as upregulated with age and that can be stimulated by an increased ECM stiffness. Another consequence of higher synthesis of UDP-GlcNAc is increased protein O-GlcNAcylation which has been implicated in sensing cellular stressors, cell-cycle alterations, transcription, protein turnover, calcium handling, bioenergetics, and nutrient levels [123] associated with a variety of heart diseases (e.g., hypertrophic heart, ischemia, and heart failure) [124–126]. Proteins involved in cardiac hypertrophic growth are among the targets of O-GlcNAcylation, including NFAT whose activity is O-GlcNAcylation-dependent [127]. Therefore, metabolism-mediated changes in gene expression, metabolite signaling, and the channeling of glucose-derived carbons toward anabolic pathways seem critical for the age-related GAG accumulation, ECM stiffness, and intracellular signaling modulation that culminate in the pathological hypertrophy of the heart.

The work reported here has several unique features. By using a NHP model across the adult lifespan we can efficiently recapitulate the aging effect on the heart’s LV in similar individuals, living in the same conditions, being exposed to the same environment, and eating a healthy low-fat and low-simple sugar diet. These animals live an optimal healthy life. In contrast, the available human heart transcriptomics data have failed to provide significant insights into cardiac healthy aging, likely due to the interplay of multiple variables encompassing environmental factors, and the challenges associated with tissue collection, including post-mortem collection times. This highlights the significance of investigating healthy heart tissue collected using rigorously defined and standardized protocols, thereby increasing the relevance of this study. While some aspects of cardiac aging in NHP exhibit a degree of independence from sex [18], which is a limitation of our study, it is imperative to acknowledge that this investigation was conducted exclusively on the understudied female subjects. Consequently, future studies of a similar nature encompassing male subjects are required to ascertain whether the mechanisms elucidated herein maintain their prominence or manifest sex-specific variations. While this is the largest reported study of NHP, and we have applied robust statistical methodology involving longitudinal regression, the inclusion of more animals in future validation studies will likely offer additional insights into the temporal sequence of the identified mechanisms in a large number of animals and further help in preventing cardiac stiffness- and hypertrophy-derived cardiac dysfunction. Changes in the many different cell types in the heart and their individual contributions to the overall GAG synthetic machinery and GAG content in the heart can be relevant in future studies, as fibroblasts and myofibroblasts are responsible for a large proportion of the GAG/ECM production. The identification of the temporal order of events during heart aging that culminate in CVD is critical to improving early diagnosis. Moreover, preventing cardiac metabolic remodeling and GAG accumulation may result in therapeutic approaches to prevent age-related CVD.

## Acknowledgments

This research was funded by the ERDF funds through the Operational Programme for Competitiveness–COMPETE 2020 and national funds by Foundation for Science and Technology under FCT-Post-doctoral Fellowship (SPP, SFRH/BPD/116061/2016), FCT-doctoral Fellowship (LFG, SFRH/BD/5539/2020) project grant CENTRO-01-0246-FEDER-000010 (Multidisciplinary Institute of Ageing in Coimbra), strategic projects UIDB/04539/2020, UIDP/04539/2020, LA/P/0058/2020). National Institutes of Health (NIH) grants U19 AG057758 and P51 OD011133. It was also funded by the European Union (HORIZON-HLTH-2022-STAYHLTH-101080329). Views and opinions expressed are however these of the author(s) only and do not necessary reflect those of the European Union or the Health and Digital Executive Agency. Neither the European Union nor the granting authority can be held responsible for them. The funding agencies had no role in study design, data collection and analysis, the decision to publish, or the preparation of this document. There are no conflicts of interest associated with this work.

## Data Availability Statement

The data supporting the findings of this study are openly available in the repository Gene Expression Omnibus (GEO; http://www.ncbi.nlm.nih.gov/geo/)—GEO Series accession number GSE245036.

---

**Figure S1.**
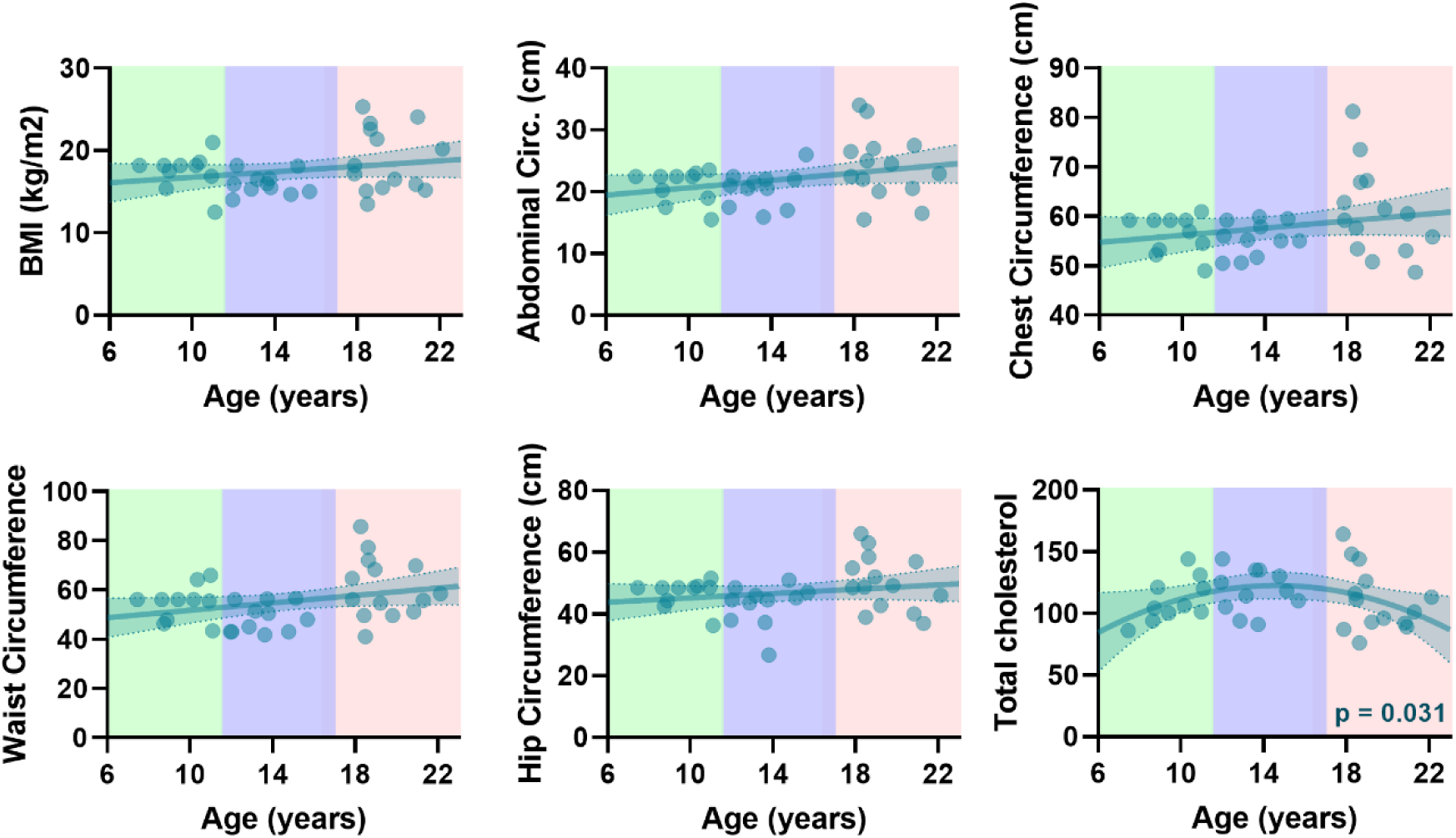
Impact of aging on baboon morphometrics and blood clinical data. Variation of morphometric (BMI, abdominal, chest, waist, and hip circumference) and blood clinical (total cholesterol) measurements with age. Circ. – circumference.

**Figure S2.**
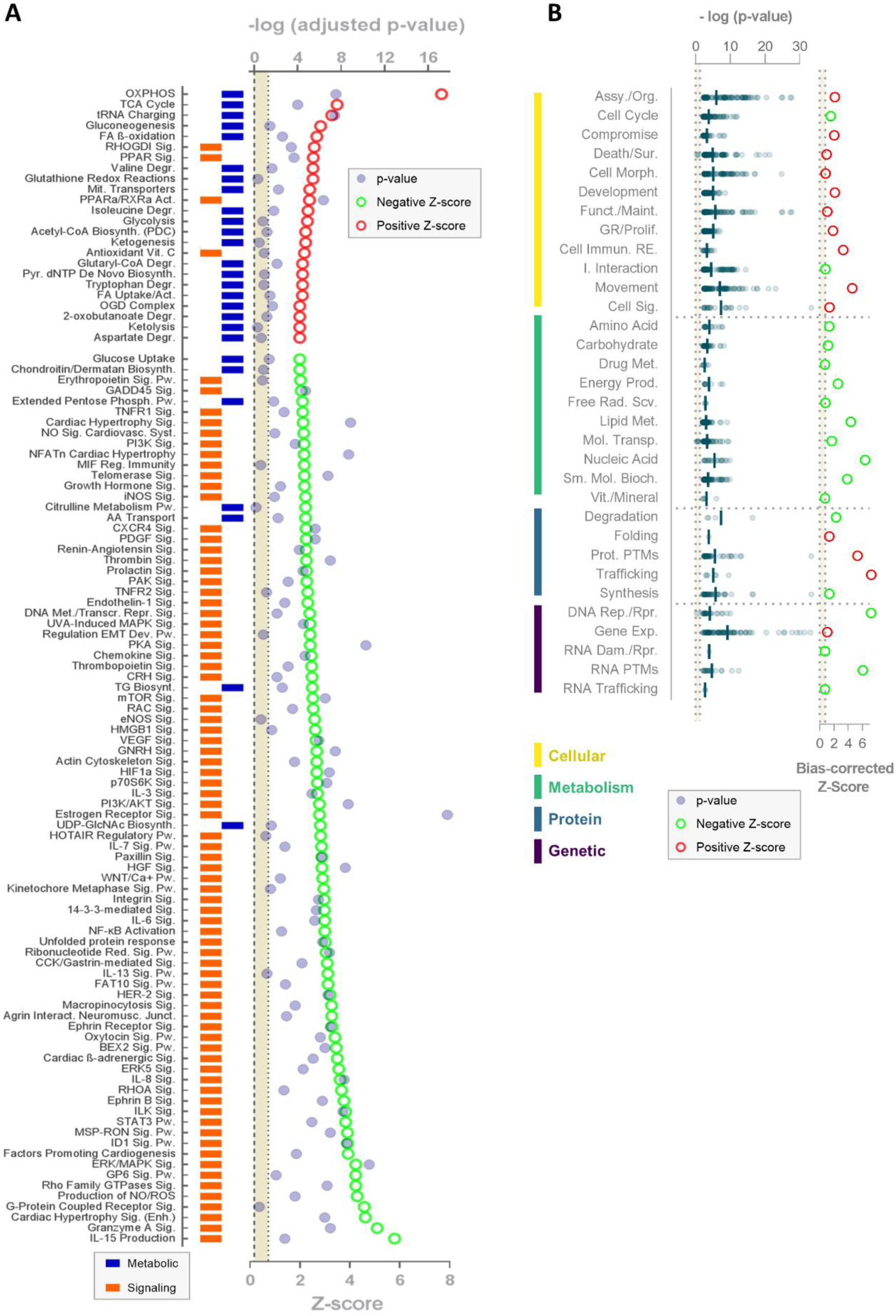
Cardiac age-related transcriptomic pathway enrichment analysis in NHP females. (A) Comparison of canonical pathways predicted activation and inhibition (absolute z-score≥2) with age. Positive z-scores are represented in red, negative z-scores in green, and the p-value in blue. The color depicts metabolic (blue) or signaling (orange) canonical pathways. (B) Biofunctions identified for the pathway enrichment analysis. Each dot represents the -log(p-value) of a biofunction. The color represents cellular (in yellow), metabolic (in green), protein-related (in blue), and genomic (in purple) biofunctions. Each circle represents the bias-corrected z-score and the respective color a negative, in green, or a positive, in red, directionality. Act. – activation; Assy. – assembly; Bioch. – biochemical; Biosynth. – biosynthesis; Dam. – damage; Degr. – degradation; Enh. – enhanced; Exp. – expression; FA – fatty acid; Funct. – function; GR – growth; Immun. – immunity; Interact. – interaction; Junct. – junction; Maint. – maintenance; Met. – metabolism; Met. – methylation; Mol. – molecular; Morph. – morphology; Neuromus. – neuromuscular; Org. – organization; OXPHOS – oxidative phosphorylation; Phosph. – phosphate; Prod. – production; Prolif. – proliferation; Prot. PTMs – protein post-translational modifications; Pw. – pathway; Rad. – radical; RE. – response; Red. – reductase; Reg. – regulatory; Rep. – replication; Repr. – Repression; RNA PTMs – RNA post-transcriptional modifications; Rpr. – repair; Scv. – scavenger; Sig. – signaling; Sm. – small; Sur. – survival; Syst. – system; TCA - tricarboxylic acid; Transcr. – transcriptional; Transp. – transport; Vit. – vitamin.

**Figure S3.**
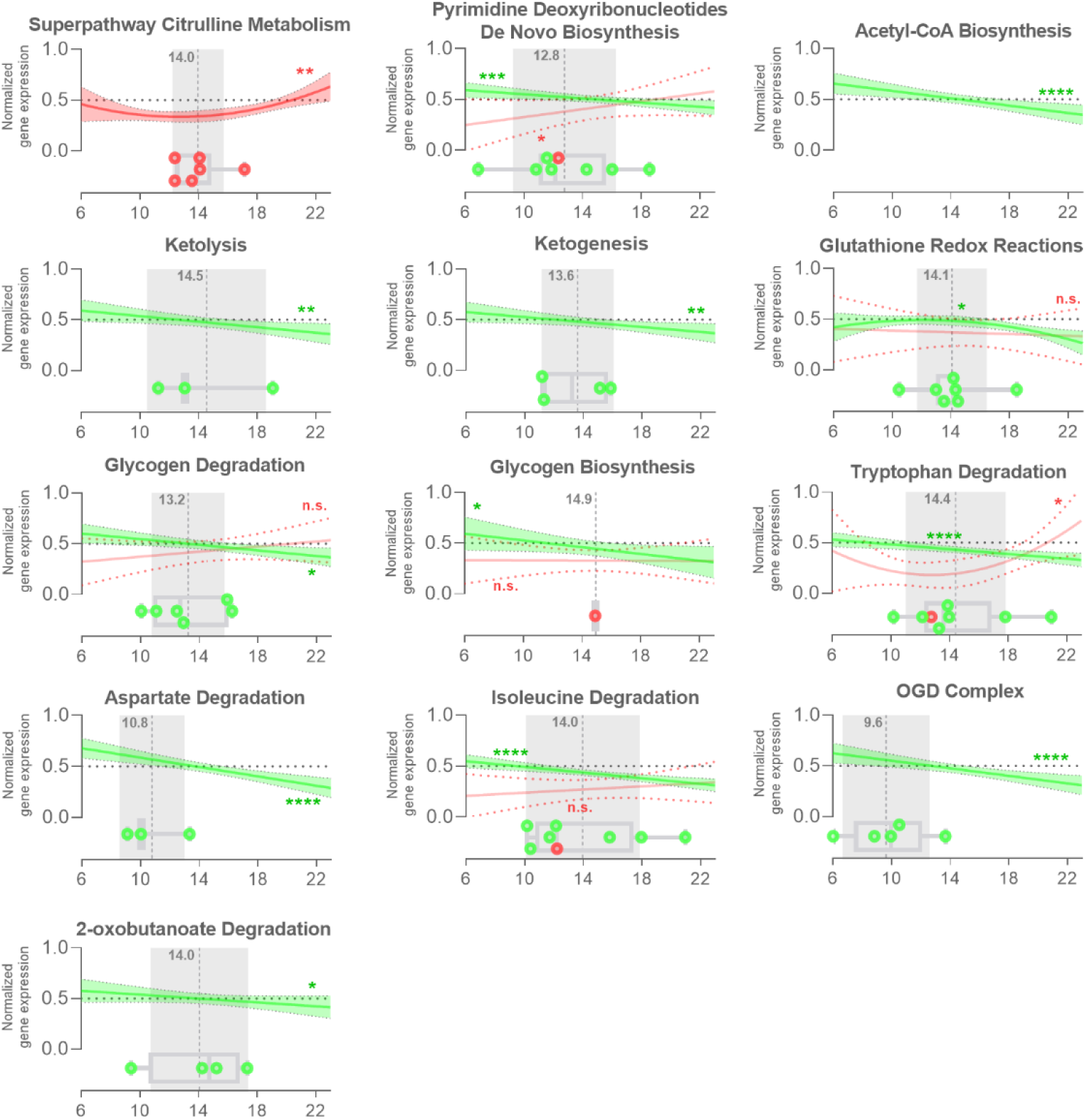
Longitudinal gene expression of pathways involved in metabolism in the NHP female heart. Longitudinal profile of relative transcript expression with age, boxplot with transcripts tipping points (red – upregulated, green – downregulated), and the age of the average tipping point (grey) from the metabolic canonical pathway.

## Bibliography

[1] Majnaric LT, Bosnic Z, Kurevija T, Wittlinger T. Cardiovascular risk and aging: the need for a more comprehensive understanding. J Geriatr Cardiol 2021;18:462. 10.11909/J.ISSN.1671-5411.2021.06.004.

[2] Obas V, Vasan RS. The aging heart. Clin Sci 2018;132:1367–82. 10.1042/CS20171156.

[3] Wang X, Lu Y, Xie Y, Shen J, Xiang M. Emerging roles of proteoglycans in cardiac remodeling. Int J Cardiol 2019;278:192–8. 10.1016/j.ijcard.2018.11.125.

[4] Benjamin EJ, Blaha MJ, Chiuve SE, Cushman M, Das SR, Deo R, et al. Heart Disease and Stroke Statistics-2017 Update: A Report From the American Heart Association. Circulation 2017;135:e146–603. 10.1161/CIR.0000000000000485.

[5] Timmis A, Townsend N, Gale CP, Torbica A, Lettino M, Petersen SE, et al. European Society of Cardiology: Cardiovascular Disease Statistics 2019. Eur Heart J 2020;41:12–85. 10.1093/EURHEARTJ/EHZ859.

[6] Taegtmeyer H, Young ME, Lopaschuk GD, Abel ED, Brunengraber H, Darley-Usmar V, et al. Assessing Cardiac Metabolism. vol. 118. 2016. 10.1161/RES.0000000000000097.

[7] Ritterhoff J, Tian R. Metabolism in cardiomyopathy: every substrate matters. Cardiovasc Res 2017;113:411–21. 10.1093/CVR/CVX017.

[8] Umbarawan Y, Syamsunarno MRAA, Koitabashi N, Obinata H, Yamaguchi A, Hanaoka H, et al. Myocardial fatty acid uptake through CD36 is indispensable for sufficient bioenergetic metabolism to prevent progression of pressure overload-induced heart failure. Sci Reports 2018 81 2018;8:1–13. 10.1038/s41598-018-30616-1.

[9] Riera CE, Dillin A. Tipping the metabolic scales towards increased longevity in mammals. Nat Cell Biol 2015 173 2015;17:196–203. 10.1038/ncb3107.

[10] Bodyak N, Kang PM, Hiromura M, Sulijoadikusumo I, Horikoshi N, Khrapko K, et al. Gene expression profiling of the aging mouse cardiac myocytes. Nucleic Acids Res 2002;30:3788–94. 10.1093/NAR/GKF497.

[11] Ma Y, Li J. Metabolic Shifts during Aging and Pathology. Compr Physiol 2015;5:667. 10.1002/CPHY.C140041.

[12] Lee CK, Allison DB, Brand J, Weindruch R, Prolla TA. Transcriptional profiles associated with aging and middle age-onset caloric restriction in mouse hearts. Proc Natl Acad Sci U S A 2002;99:14988–93. 10.1073/PNAS.232308999.

[13] Tran DH, May HI, Li Q, Luo X, Huang J, Zhang G, et al. Chronic activation of hexosamine biosynthesis in the heart triggers pathological cardiac remodeling. Nat Commun 2020 111 2020;11:1–15. 10.1038/s41467-020-15640-y.

[14] Ritterhoff J, Young S, Villet O, Shao D, Carnevale Neto F, Bettcher LF, et al. Metabolic Remodeling Promotes Cardiac Hypertrophy by Directing Glucose to Aspartate Biosynthesis. Circ Res 2020;126:182–96. 10.1161/CIRCRESAHA.119.315483.

[15] Gibb AA, Hill BG. Metabolic coordination of physiological and pathological cardiac remodeling. Circ Res 2018;123:107–28. 10.1161/CIRCRESAHA.118.312017.

[16] Tracy E, Rowe G, LeBlanc AJ. Tissue Remodeling: From Regeneration to Fibrosis: Cardiac tissue remodeling in healthy aging: the road to pathology. Am J Physiol - Cell Physiol 2020;319:C166. 10.1152/AJPCELL.00021.2020.

[17] Kuo AH, Li C, Mattern V, Huber HF, Comuzzie A, Cox L, et al. Sex-Dimorphic Acceleration of Pericardial, Subcutaneous, and Plasma Lipid Increase in Offspring of Poorly Nourished Baboons. Int J Obes (Lond) 2018;42:1092. 10.1038/S41366-018-0008-2.

[18] Kuo AH, Li C, Huber HF, Nathanielsz PW, Clarke GD. Ageing changes in biventricular cardiac function in male and female baboons (Papio spp.). J Physiol 2018;596:5083. 10.1113/JP276338.

[19] Kuo AH, Li C, Li J, Huber HF, Nathanielsz PW, Clarke GD. Cardiac remodelling in a baboon model of intrauterine growth restriction mimics accelerated ageing. J Physiol 2017;595:1093. 10.1113/JP272908.

[20] Kuo AH, Li C, Huber HF, Schwab M, Nathanielsz PW, Clarke GD. Maternal nutrient restriction during pregnancy and lactation leads to impaired right ventricular function in young adult baboons. J Physiol 2017;595:4245. 10.1113/JP273928.

[21] Zhao R-R, Ackers-Johnson M, Stenzig J, Chen C, Ding T, Zhou Y, et al. Targeting Chondroitin Sulfate Glycosaminoglycans to Treat Cardiac Fibrosis in Pathological Remodeling. Circulation 2018;137:2497–513. 10.1161/CIRCULATIONAHA.117.030353.

[22] Luong H, Singh S, Patil M, Krishnamurthy P. Cardiac glycosaminoglycans and structural alterations during chronic stress-induced depression-like behavior in mice. Am J Physiol - Hear Circ Physiol 2021;320:H2044–57. 10.1152/AJPHEART.00635.2020/ASSET/IMAGES/LARGE/AJPHEART.00635.2020_F008.JPEG.

[23] Rienks M, Papageorgiou AP, Frangogiannis NG, Heymans S. Myocardial Extracellular Matrix. Circ Res 2014;114:872–88. 10.1161/CIRCRESAHA.114.302533.

[24] Soares Da Costa D, Reis RL, Pashkuleva I. Sulfation of Glycosaminoglycans and Its Implications in Human Health and Disorders. Https://DoiOrg/101146/Annurev-Bioeng-071516-044610 2017;19:1–26. 10.1146/ANNUREV-BIOENG-071516-044610.

[25] Mizumoto S, Yamada S. Congenital Disorders of Deficiency in Glycosaminoglycan Biosynthesis. Front Genet 2021;12:1632. 10.3389/FGENE.2021.717535/BIBTEX.

[26] Jana S, Zhang H, Lopaschuk GD, Freed DH, Sergi C, Kantor PF, et al. Disparate remodeling of the extracellular matrix and proteoglycans in failing pediatric versus adult hearts. J Am Heart Assoc 2018;7. 10.1161/JAHA.118.010427.

[27] Bartling B, Niemann K, Pliquett RU, Treede H, Simm A. Altered gene expression pattern indicates the differential regulation of the immune response system as an important factor in cardiac aging. Exp Gerontol 2019;117:13–20. 10.1016/J.EXGER.2018.05.001.

[28] De Majo F, Hegenbarth J-C, Rühle F, Bär C, Thum T, de Boer M, et al. Dichotomy between the transcriptomic landscape of naturally versus accelerated aged murine hearts. Sci Rep 2020;10:8136. 10.1038/s41598-020-65115-9.

[29] Haghighi K, Kolokathis F, Pater L, Lynch RA, Asahi M, Gramolini AO, et al. Human phospholamban null results in lethal dilated cardiomyopathy revealing a critical difference between mouse and human. J Clin Invest 2003;111:869–76. 10.1172/JCI17892.

[30] Greenig M, Melville A, Huntley D, Isalan M, Mielcarek M. Cross-Sectional Transcriptional Analysis of the Aging Murine Heart. Front Mol Biosci 2020;7. 10.3389/FMOLB.2020.565530.

[31] Cox L, Olivier M. SIRT2 counteracts primate cardiac aging. Nat Aging 2023 310 2023;3:1178–9. 10.1038/s43587-023-00496-w.

[32] Peterson LR, Soto PF, Herrero P, Schechtman KB, Dence C, Gropler RJ. Sex differences in myocardial oxygen and glucose metabolism. J Nucl Cardiol 2007 144 2007;14:573–81. 10.1016/J.NUCLCARD.2007.03.001.

[33] Peterson LR, Soto PF, Herrero P, Mohammed BS, Avidan MS, Schechtman KB, et al. Impact of Gender on the Myocardial Metabolic Response to Obesity. JACC Cardiovasc Imaging 2008;1:424–33. 10.1016/J.JCMG.2008.05.004.

[34] Bronikowski AM, Alberts SC, Altmann J, Packer C, Dee Carey K, Tatar M. The aging baboon: Comparative demography in a non-human primate. Proc Natl Acad Sci U S A 2002;99:9591–5. 10.1073/PNAS.142675599/SUPPL_FILE/6755TABLE2.PDF.

[35] Schlabritz-Loutsevitch N, Ballesteros B, Dudley C, Jenkins S, Hubbard G, Burton GJ, et al. Moderate maternal nutrient restriction, but not glucocorticoid administration, leads to placental morphological changes in the baboon (Papio sp.). Placenta 2007;28:783–93. 10.1016/J.PLACENTA.2006.11.012.

[36] Yang S, Gerow KG, Huber HF, Considine MM, Li C, Mattern V, et al. A decline in female baboon hypothalamo-pituitary-adrenal axis activity anticipates aging. Aging (Albany NY) 2017;9:1375–85. 10.18632/AGING.101235.

[37] Chavez AO, Gastaldelli A, Guardado-Mendoza R, Lopez-Alvarenga JC, Michelle MM, Elizabeth ME, et al. Predictive models of insulin resistance derived from simple morphometric and biochemical indices related to obesity and the metabolic syndrome in baboons. Cardiovasc Diabetol 2009;8:22. 10.1186/1475-2840-8-22.

[38] Li C, Jenkins S, Mattern V, Comuzzie AG, Cox LA, Huber HF, et al. Effect of moderate, 30 percent global maternal nutrient reduction on fetal and postnatal baboon phenotype. J Med Primatol 2017;46:293–303. 10.1111/JMP.12290.

[39] Kim D, Langmead B, Salzberg SL. HISAT: a fast spliced aligner with low memory requirements. Nat Methods 2015 124 2015;12:357–60. 10.1038/nmeth.3317.

[40] Xing Y, Yu T, Wu YN, Roy M, Kim J, Lee C. An expectation-maximization algorithm for probabilistic reconstructions of full-length isoforms from splice graphs. Nucleic Acids Res 2006;34:3150–60. 10.1093/NAR/GKL396.

[41] Robinson MD, Oshlack A. A scaling normalization method for differential expression analysis of RNA-seq data. Genome Biol 2010;11. 10.1186/GB-2010-11-3-R25.

[42] Langfelder P, Horvath S. WGCNA: an R package for weighted correlation network analysis. BMC Bioinformatics 2008;9:559. 10.1186/1471-2105-9-559.

[43] Wang W, Shi L, Qin Y, Li F. Research and Application of Chondroitin Sulfate/Dermatan Sulfate-Degrading Enzymes. Front Cell Dev Biol 2020;8:1435. 10.3389/FCELL.2020.560442/BIBTEX.

[44] Baranyi J, Pin C. Estimating Bacterial Growth Parameters by Means of Detection Times. Appl Environ Microbiol 1999;65:732. 10.1128/AEM.65.2.732-736.1999.

[45] Baudouy D, Michiels JF, Vukolic A, Wagner KD, Wagner N. Echocardiographic and Histological Examination of Cardiac Morphology in the Mouse. J Vis Exp 2017;2017:55843. 10.3791/55843.

[46] da Silva FS, Aquino de Souza NCS, de Moraes MV, Abreu BJ, de Oliveira MF. CmyoSize: An ImageJ macro for automated analysis of cardiomyocyte size in images of routine histology staining. Ann Anat 2022;241. 10.1016/J.AANAT.2022.151892.

[47] Demšar J, Curk T, Erjavec A, Gorup Č, Hočevar T, Milutinovič M, et al. Orange: Data Mining Toolbox in Python. J Mach Learn Res 2013;14:2349–53.

[48] Lesnefsky EJ, Chen Q, Hoppel CL. Mitochondrial Metabolism in Aging Heart. Circ Res 2016;118:1593–611. 10.1161/CIRCRESAHA.116.307505.

[49] Ge H, Tian M, Pei Q, Tan F, Pei H. Extracellular Matrix Stiffness: New Areas Affecting Cell Metabolism. Front Oncol 2021;11:8. 10.3389/FONC.2021.631991/BIBTEX.

[50] Sciarretta S, Volpe M, Sadoshima J. Mammalian target of rapamycin signaling in cardiac physiology and disease. Circ Res 2014;114:549–64. 10.1161/CIRCRESAHA.114.302022.

[51] Ritchie H, Roser M. Age Structure. Publ Online OurWorldInDataOrg 2019. https://ourworldindata.org/age-structure (accessed October 24, 2022).

[52] Conrad N, Judge A, Tran J, Mohseni H, Hedgecott D, Crespillo AP, et al. Temporal trends and patterns in heart failure incidence: a population-based study of 4 million individuals. Lancet 2018;391:572–80. 10.1016/S0140-6736(17)32520-5.

[53] Mora S, Dugani SB, Moorthy MV, Li C, Demler O V., Alsheikh-Ali AA, et al. Association of Lipid, Inflammatory, and Metabolic Biomarkers With Age at Onset for Incident Coronary Heart Disease in Women. JAMA Cardiol 2021;6:1. 10.1001/JAMACARDIO.2020.7073.

[54] Foley DL, Mackinnon A, Watts GF, Shaw JE, Magliano DJ, Castle DJ, et al. Cardiometabolic Risk Indicators That Distinguish Adults with Psychosis from the General Population, by Age and Gender. PLoS One 2013;8:e82606. 10.1371/JOURNAL.PONE.0082606.

[55] Yi SW, Park S, Lee YH, Park HJ, Balkau B, Yi JJ. Association between fasting glucose and all-cause mortality according to sex and age: a prospective cohort study. Sci Reports 2017 71 2017;7:1–9. 10.1038/s41598-017-08498-6.

[56] Feng L, Nian S, Tong Z, Zhu Y, Li Y, Zhang C, et al. Age-related trends in lipid levels: a large-scale cross-sectional study of the general Chinese population. BMJ Open 2020;10:e034226. 10.1136/BMJOPEN-2019-034226.

[57] Yeung KR, Chiu CL, Pears S, Heffernan SJ, Makris A, Hennessy A, et al. A Cross-Sectional Study of Ageing and Cardiovascular Function over the Baboon Lifespan. PLoS One 2016;11. 10.1371/JOURNAL.PONE.0159576.

[58] Silva AC, Pereira C, Fonseca ACRG, Pinto-do-Ó P, Nascimento DS. Bearing My Heart: The Role of Extracellular Matrix on Cardiac Development, Homeostasis, and Injury Response. Front Cell Dev Biol 2021;8:1705. 10.3389/FCELL.2020.621644/BIBTEX.

[59] López-otín C, Blasco MA, Partridge L, Serrano M, Kroemer G. The Hallmarks of Aging. Cell 2013;153:1194–217. 10.1016/j.cell.2013.05.039.The.

[60] Ye Y, Yang K, Liu H, Yu Y, Song M, Huang D, et al. SIRT2 counteracts primate cardiac aging via deacetylation of STAT3 that silences CDKN2B. Nat Aging 2023 310 2023;3:1269–87. 10.1038/s43587-023-00486-y.

[61] Loffredo FS, Steinhauser ML, Jay SM, Gannon J, Pancoast JR, Yalamanchi P, et al. Growth differentiation factor 11 is a circulating factor that reverses age-related cardiac hypertrophy. Cell 2013;153:828–39. 10.1016/J.CELL.2013.04.015.

[62] Bradshaw AD, Baicu CF, Rentz TJ, Van Laer AO, Bonnema DD, Zile MR. Age-dependent alterations in fibrillar collagen content and myocardial diastolic function: role of SPARC in post-synthetic procollagen processing. Am J Physiol Heart Circ Physiol 2010;298. 10.1152/AJPHEART.00474.2009.

[63] Annoni G, Luvarà G, Arosio B, Gagliano N, Fiordaliso F, Santambrogio D, et al. Age-dependent expression of fibrosis-related genes and collagen deposition in the rat myocardium. Mech Ageing Dev 1998;101:57–72. 10.1016/S0047-6374(97)00165-6.

[64] Liu J, Masurekar MR, Vatner DE, Jyothirmayi GN, Regan TJ, Vatner SF, et al. Glycation end-product cross-link breaker reduces collagen and improves cardiac function in aging diabetic heart. Am J Physiol Heart Circ Physiol 2003;285. 10.1152/AJPHEART.00516.2003.

[65] Gazoti Debessa CR, Mesiano Maifrino LB, Rodrigues de Souza R. Age related changes of the collagen network of the human heart. Mech Ageing Dev 2001;122:1049–58. 10.1016/S0047-6374(01)00238-X.

[66] Huynh MB, Morin C, Carpentier G, Garcia-Filipe S, Talhas-Perret S, Barbier-Chassefière V, et al. Age-related Changes in Rat Myocardium Involve Altered Capacities of Glycosaminoglycans to Potentiate Growth Factor Functions and Heparan Sulfate-altered Sulfation. J Biol Chem 2012;287:11363–73. 10.1074/JBC.M111.335901.

[67] Shi D, Sheng A, Chi L. Glycosaminoglycan-Protein Interactions and Their Roles in Human Disease. Front Mol Biosci 2021;8:64. 10.3389/FMOLB.2021.639666/BIBTEX.

[68] Tuysuz B, Mizumoto S, Sugahara K, Çelebi A, Mundlos S, Turkmen S. Omani-type spondyloepiphyseal dysplasia with cardiac involvement caused by a missense mutation in CHST3. Clin Genet 2009;75:375–83. 10.1111/J.1399-0004.2009.01167.X.

[69] Dündar M, Müller T, Zhang Q, Pan J, Steinmann B, Vodopiutz J, et al. Loss of dermatan-4-sulfotransferase 1 function results in adducted thumb-clubfoot syndrome. Am J Hum Genet 2009;85:873–82. 10.1016/J.AJHG.2009.11.010.

[70] Baasanjav S, Al-Gazali L, Hashiguchi T, Mizumoto S, Fischer B, Horn D, et al. Faulty initiation of proteoglycan synthesis causes cardiac and joint defects. Am J Hum Genet 2011;89:15–27. 10.1016/J.AJHG.2011.05.021.

[71] Munns CF, Fahiminiya S, Poudel N, Munteanu MC, Majewski J, Sillence DO, et al. Homozygosity for Frameshift Mutations in XYLT2 Result in a Spondylo-Ocular Syndrome with Bone Fragility, Cataracts, and Hearing Defects. Am J Hum Genet 2015;96:971. 10.1016/J.AJHG.2015.04.017.

[72] Veugelers M, De Cat B, Muyldermans SY, Reekmans G, Delande N, Frints S, et al. Mutational analysis of the GPC3/GPC4 glypican gene cluster on Xq26 in patients with Simpson-Golabi-Behmel syndrome: identification of loss-of-function mutations in the GPC3 gene. Hum Mol Genet 2000;9:1321–8. 10.1093/HMG/9.9.1321.

[73] Young WJ, Lahrouchi N, Isaacs A, Duong TV, Foco L, Ahmed F, et al. Genetic analyses of the electrocardiographic QT interval and its components identify additional loci and pathways. Nat Commun 2022;13. 10.1038/S41467-022-32821-Z.

[74] Pirruccello JP, Chaffin MD, Chou EL, Fleming SJ, Lin H, Nekoui M, et al. Deep learning enables genetic analysis of the human thoracic aorta. Nat Genet 2022;54:40–51. 10.1038/S41588-021-00962-4.

[75] Hoffmann TJ, Ehret GB, Nandakumar P, Ranatunga D, Schaefer C, Kwok P-Y, et al. Genome-wide association analyses using electronic health records identify new loci influencing blood pressure variation. Nat Genet 2017;49:54–64. 10.1038/ng.3715.

[76] Wain L V., Vaez A, Jansen R, Joehanes R, van der Most PJ, Erzurumluoglu AM, et al. Novel Blood Pressure Locus and Gene Discovery Using Genome-Wide Association Study and Expression Data Sets From Blood and the Kidney. Hypertension 2017;70. 10.1161/HYPERTENSIONAHA.117.09438.

[77] Sakaue S, Kanai M, Tanigawa Y, Karjalainen J, Kurki M, Koshiba S, et al. A cross-population atlas of genetic associations for 220 human phenotypes. Nat Genet 2021;53:1415–24. 10.1038/s41588-021-00931-x.

[78] Roselli C, Chaffin MD, Weng L-C, Aeschbacher S, Ahlberg G, Albert CM, et al. Multi-ethnic genome-wide association study for atrial fibrillation. Nat Genet 2018;50:1225–33. 10.1038/s41588-018-0133-9.

[79] Cárcel-Márquez J, Muiño E, Gallego-Fabrega C, Cullell N, Lledós M, Llucià-Carol L, et al. A Polygenic Risk Score Based on a Cardioembolic Stroke Multitrait Analysis Improves a Clinical Prediction Model for This Stroke Subtype. Front Cardiovasc Med 2022;9. 10.3389/fcvm.2022.940696.

[80] Nielsen JB, Thorolfsdottir RB, Fritsche LG, Zhou W, Skov MW, Graham SE, et al. Biobank-driven genomic discovery yields new insight into atrial fibrillation biology. Nat Genet 2018;50:1234–9. 10.1038/s41588-018-0171-3.

[81] Ellinor PT, Lunetta KL, Glazer NL, Pfeufer A, Alonso A, Chung MK, et al. Common variants in KCNN3 are associated with lone atrial fibrillation. Nat Genet 2010;42:240–4. 10.1038/ng.537.

[82] Takahashi Y, Yamazaki K, Kamatani Y, Kubo M, Matsuda K, Asai S. A genome-wide association study identifies a novel candidate locus at the DLGAP1 gene with susceptibility to resistant hypertension in the Japanese population. Sci Rep 2021;11:19497. 10.1038/s41598-021-98144-z.

[83] Chen J, Wang W, Li Z, Xu C, Tian X, Zhang D. Heritability and genome-wide association study of blood pressure in Chinese adult twins. Mol Genet Genomic Med 2021;9. 10.1002/mgg3.1828.

[84] Boua PR, Brandenburg J-T, Choudhury A, Sorgho H, Nonterah EA, Agongo G, et al. Genetic associations with carotid intima-media thickness link to atherosclerosis with sex-specific effects in sub-Saharan Africans. Nat Commun 2022;13:855. 10.1038/s41467-022-28276-x.

[85] Benjamins JW, Yeung MW, van de Vegte YJ, Said MA, van der Linden T, Ties D, et al. Genomic insights in ascending aortic size and distensibility. EBioMedicine 2022;75. 10.1016/J.EBIOM.2021.103783.

[86] Song Y, Choi J-E, Kwon Y-J, Chang H-J, Kim JO, Park D-H, et al. Identification of susceptibility loci for cardiovascular disease in adults with hypertension, diabetes, and dyslipidemia. J Transl Med 2021;19:85. 10.1186/s12967-021-02751-3.

[87] Justice AE, Howard AG, Chittoor G, Fernandez-Rhodes L, Graff M, Voruganti VS, et al. Genome-wide association of trajectories of systolic blood pressure change. BMC Proc 2016;10:56. 10.1186/s12919-016-0050-9.

[88] Kichaev G, Bhatia G, Loh P-R, Gazal S, Burch K, Freund MK, et al. Leveraging Polygenic Functional Enrichment to Improve GWAS Power. Am J Hum Genet 2019;104:65–75. 10.1016/j.ajhg.2018.11.008.

[89] Gouveia MH, Bentley AR, Leonard H, Meeks KAC, Ekoru K, Chen G, et al. Trans-ethnic meta-analysis identifies new loci associated with longitudinal blood pressure traits. Sci Rep 2021;11:4075. 10.1038/s41598-021-83450-3.

[90] Drago A, Kure Fischer E. A molecular pathway analysis informs the genetic risk for arrhythmias during antipsychotic treatment. Int Clin Psychopharmacol 2018;33:1–14. 10.1097/YIC.0000000000000198.

[91] Francis CM, Futschik ME, Huang J, Bai W, Sargurupremraj M, Teumer A, et al. Genome-wide associations of aortic distensibility suggest causality for aortic aneurysms and brain white matter hyperintensities. Nat Commun 2022;13. 10.1038/S41467-022-32219-X.

[92] Tcheandjieu C, Xiao K, Tejeda H, Lynch JA, Ruotsalainen S, Bellomo T, et al. High heritability of ascending aortic diameter and trans-ancestry prediction of thoracic aortic disease. Nat Genet 2022;54:772–82. 10.1038/S41588-022-01070-7.

[93] de las Fuentes L, Sung YJ, Noordam R, Winkler T, Feitosa MF, Schwander K, et al. Gene-educational attainment interactions in a multi-ancestry genome-wide meta-analysis identify novel blood pressure loci. Mol Psychiatry 2021;26:2111–25. 10.1038/S41380-020-0719-3.

[94] van der Harst P, Verweij N. Identification of 64 Novel Genetic Loci Provides an Expanded View on the Genetic Architecture of Coronary Artery Disease. Circ Res 2018;122:433–43. 10.1161/CIRCRESAHA.117.312086.

[95] Kumar A, Chauhan G, Sharma S, Dabla S, Sylaja PN, Chaudhary N, et al. Association of SUMOylation Pathway Genes With Stroke in a Genome-Wide Association Study in India. Neurology 2021;97:e345–56. 10.1212/WNL.0000000000012258.

[96] Li C, He J, Chen J, Zhao J, Gu D, Hixson JE, et al. Genome-Wide Gene–Sodium Interaction Analyses on Blood Pressure. Hypertension 2016;68:348–55. 10.1161/HYPERTENSIONAHA.115.06765.

[97] Ehret GB, Ferreira T, Chasman DI, Jackson AU, Schmidt EM, Johnson T, et al. The genetics of blood pressure regulation and its target organs from association studies in 342,415 individuals. Nat Genet 2016;48:1171–84. 10.1038/ng.3667.

[98] Vistnes M, Aronsen JM, Lunde IG, Sjaastad I, Carlson CR, Christensen G. Pentosan Polysulfate Decreases Myocardial Expression of the Extracellular Matrix Enzyme ADAMTS4 and Improves Cardiac Function In Vivo in Rats Subjected to Pressure Overload by Aortic Banding. PLoS One 2014;9:e89621. 10.1371/JOURNAL.PONE.0089621.

[99] Waehre A, Halvorsen B, Yndestad A, Husberg C, Sjaastad I, Nygård S, et al. Lack of Chemokine Signaling through CXCR5 Causes Increased Mortality, Ventricular Dilatation and Deranged Matrix during Cardiac Pressure Overload. PLoS One 2011;6:e18668. 10.1371/JOURNAL.PONE.0018668.

[100] Engebretsen KVT, Waehre A, Bjørnstad JL, Skrbic B, Sjaastad I, Behmen D, et al. Decorin, lumican, and their GAG chain-synthesizing enzymes are regulated in myocardial remodeling and reverse remodeling in the mouse. J Appl Physiol 2013;114:988–97. 10.1152/JAPPLPHYSIOL.00793.2012/ASSET/IMAGES/LARGE/ZDG008130 5390007.JPEG.

[101] Engebretsen KVT, Lunde IG, Strand ME, Waehre A, Sjaastad I, Marstein HS, et al. Lumican is increased in experimental and clinical heart failure, and its production by cardiac fibroblasts is induced by mechanical and proinflammatory stimuli. FEBS J 2013;280:2382–98. 10.1111/FEBS.12235.

[102] Chen SW, Tung YC, Jung SM, Chu Y, Lin PJ, Kao WWY, et al. Lumican-null mice are susceptible to aging and isoproterenol-induced myocardial fibrosis. Biochem Biophys Res Commun 2017;482:1304–11. 10.1016/J.BBRC.2016.12.033.

[103] del Monte-Nieto G, Fischer JW, Gorski DJ, Harvey RP, Kovacic JC. Basic Biology of Extracellular Matrix in the Cardiovascular System, Part 1/4: JACC Focus Seminar. J Am Coll Cardiol 2020;75:2169–88. 10.1016/J.JACC.2020.03.024.

[104] Grandoch M, Kohlmorgen C, Melchior-Becker A, Feldmann K, Homann S, Müller J, et al. Loss of Biglycan Enhances Thrombin Generation in Apolipoprotein E-Deficient Mice: Implications for Inflammation and Atherosclerosis. Arterioscler Thromb Vasc Biol 2016;36:e41–50. 10.1161/ATVBAHA.115.306973.

[105] Zhang L, Beeler DL, Lawrence R, Lech M, Liu J, Davis JC, et al. 6-O-Sulfotransferase-1 Represents a Critical Enzyme in the Anticoagulant Heparan Sulfate Biosynthetic Pathway. J Biol Chem 2001;276:42311–21. 10.1074/JBC.M101441200.

[106] Cai Y, Zhang X, Shen J, Jiang B, Hu D, Zhao M. Heparin-Binding Protein: A Novel Biomarker Linking Four Different Cardiovascular Diseases. Cardiol Res Pract 2020;2020. 10.1155/2020/9575373.

[107] Christensen G, Herum KM, Lunde IG. Sweet, yet underappreciated: Proteoglycans and extracellular matrix remodeling in heart disease. Matrix Biol 2019;75–76:286–99. 10.1016/J.MATBIO.2018.01.001.

[108] Vigetti D, Deleonibus S, Moretto P, Karousou E, Viola M, Bartolini B, et al. Role of UDP-N-Acetylglucosamine (GlcNAc) and O-GlcNAcylation of Hyaluronan Synthase 2 in the Control of Chondroitin Sulfate and Hyaluronan Synthesis. J Biol Chem 2012;287:35544–55. 10.1074/JBC.M112.402347.

[109] Chowdhury B, Xiang B, Liu M, Hemming R, Dolinsky VW, Triggs-Raine B. Hyaluronidase 2 Deficiency Causes Increased Mesenchymal Cells, Congenital Heart Defects, and Heart Failure. Circ Cardiovasc Genet 2017;10. 10.1161/CIRCGENETICS.116.001598.

[110] Camenisch TD, Spicer AP, Brehm-Gibson T, Biesterfeldt J, Augustine M Lou, Calabro A, et al. Disruption of hyaluronan synthase-2 abrogates normal cardiac morphogenesis and hyaluronan-mediated transformation of epithelium to mesenchyme. J Clin Invest 2000;106:349–60. 10.1172/JCI10272.

[111] De la Motte CA, Hascall VC, Drazba J, Bandyopadhyay SK, Strong SA. Mononuclear Leukocytes Bind to Specific Hyaluronan Structures on Colon Mucosal Smooth Muscle Cells Treated with Polyinosinic Acid:Polycytidylic Acid: Inter-α-Trypsin Inhibitor Is Crucial to Structure and Function. Am J Pathol 2003;163:121–33. 10.1016/S0002-9440(10)63636-X.

[112] Herum KM, Lunde IG, Skrbic B, Louch WE, Hasic A, Boye S, et al. Syndecan-4 is a key determinant of collagen cross-linking and passive myocardial stiffness in the pressure-overloaded heart. Cardiovasc Res 2015;106:217–26. 10.1093/CVR/CVV002.

[113] Hajduch E, Alessi DR, Hemmings BA, Hundal HS. Constitutive activation of protein kinase B alpha by membrane targeting promotes glucose and system A amino acid transport, protein synthesis, and inactivation of glycogen synthase kinase 3 in L6 muscle cells. Diabetes 1998;47:1006–13. 10.2337/DIABETES.47.7.1006.

[114] Barthel A, Okino ST, Liao J, Nakatani K, Li J, Whitlock JP, et al. Regulation of GLUT1 gene transcription by the serine/threonine kinase Akt1. J Biol Chem 1999;274:20281–6. 10.1074/JBC.274.29.20281.

[115] Bogatikov E, Munoz C, Pakladok T, Alesutan I, Shojaiefard M, Seebohm G, et al. Up-regulation of amino acid transporter SLC6A19 activity and surface protein abundance by PKB/Akt and PIKfyve. Cell Physiol Biochem 2012;30:1538–46. 10.1159/000343341.

[116] Hu J, Zhu XH, Ye L, Hong W, Yao KN, Chen ZY. Ursodeoxycholic acid ameliorates hepatic lipid metabolism in LO2 cells by regulating the AKT/mTOR/SREBP-1 signaling pathway. World J Gastroenterol 2019;25:1492–501. 10.3748/WJG.V25.I12.1492.

[117] Park JS, Burckhardt CJ, Lazcano R, Solis LM, Isogai T, Li L, et al. Mechanical regulation of glycolysis via cytoskeleton architecture. Nat 2020 5787796 2020;578:621–6. 10.1038/s41586-020-1998-1.

[118] Hu H, Juvekar A, Lyssiotis CA, Lien EC, Albeck JG, Oh D, et al. Phosphoinositide 3-Kinase Regulates Glycolysis through Mobilization of Aldolase from the Actin Cytoskeleton. Cell 2016;164:433–46. 10.1016/J.CELL.2015.12.042.

[119] Fernie AR, Zhang Y, Sampathkumar A. Cytoskeleton Architecture Regulates Glycolysis Coupling Cellular Metabolism to Mechanical Cues. Trends Biochem Sci 2020;45:637–8. 10.1016/J.TIBS.2020.04.003.

[120] Grilo LF, Tocantins C, Diniz MS, Gomes RM, Oliveira PJ, Matafome P, et al. Metabolic Disease Programming: From Mitochondria to Epigenetics, Glucocorticoid Signalling and Beyond. Eur J Clin Invest 2021. 10.1111/eci.13625.

[121] Tran DH, Wang Z V. Glucose Metabolism in Cardiac Hypertrophy and Heart Failure. J Am Heart Assoc 2019;8. 10.1161/JAHA.119.012673.

[122] Laczy B, Fülöp N, Onay-Besikci A, des Rosiers C, Chatham JC. Acute Regulation of Cardiac Metabolism by the Hexosamine Biosynthesis Pathway and Protein O-GlcNAcylation. PLoS One 2011;6. 10.1371/JOURNAL.PONE.0018417.

[123] Wright JLN, Collins HE, Wende AR, Chatham JC. O-GlcNAcylation and cardiovascular disease. Biochem Soc Trans 2017;45:545–53. 10.1042/BST20160164.

[124] Ledee D, Smith L, Bruce M, Kajimoto M, Isern N, Portman MA, et al. c-Myc Alters Substrate Utilization and O-GlcNAc Protein Posttranslational Modifications without Altering Cardiac Function during Early Aortic Constriction. PLoS One 2015;10:e0135262. 10.1371/JOURNAL.PONE.0135262.

[125] Cannon M V, Silljé HH, Sijbesma JW, Vreeswijk-Baudoin I, Ciapaite J, van der Sluis B, et al. Cardiac LXRα protects against pathological cardiac hypertrophy and dysfunction by enhancing glucose uptake and utilization. EMBO Mol Med 2015;7:1229–43. 10.15252/EMMM.201404669.

[126] Lunde IG, Aronsen JM, Kvaløy H, Qvigstad E, Sjaastad I, Tønnessen T, et al. Cardiac O-GlcNAc signaling is increased in hypertrophy and heart failure. Physiol Genomics 2012;44:162–72. 10.1152/PHYSIOLGENOMICS.00016.2011/SUPPL_FILE/SUPPMAT.PDF.

[127] Facundo HT, Brainard RE, Watson LJ, Ngoh GA, Hamid T, Prabhu SD, et al. O-GlcNAc signaling is essential for NFAT-mediated transcriptional reprogramming during cardiomyocyte hypertrophy. Am J Physiol Heart Circ Physiol 2012;302. 10.1152/AJPHEART.00775.2011.

